# Insular opioid signaling and distinct claustral ensembles in hedonic feeding

**DOI:** 10.1101/2024.06.10.598168

**Authors:** Jingyi Chen, Leandra Mangieri, David Marcus, Madison Martin, Veronica Porubsky, Sean Piantadosi, Sophia Mar, Phoenix Davis, Arshia Kumar, Berkan Mertan, Kaspar Podgorski, Larry Zweifel, Benjamin Land, Michael R. Bruchas

## Abstract

Stress triggers eating that is driven by palatability rather than energy need, but how stress-related signals act on the circuits that control hedonic feeding is unclear. The claustrum, a thin sheet of neurons reciprocally connected with much of the cortex, is enriched in κ-opioid receptors (KORs), the receptors for the stress-related peptide dynorphin. Here we show that a single forced swim increases palatable-food intake in mice without food restriction. Deleting or blocking claustral KOR attenuated this increase. A KOR-based sensor reported a fall in claustral opioid tone at the onset of eating. Prodynorphin (Pdyn)-expressing neurons in the anterior insular cortex (aIC) projected to the claustrum and recruited both vGAT and CaMKII-expressing claustral neurons. Single-cell imaging showed that eating engages claustral populations in distinct ways: nearly all vGAT neurons were activated, whereas KOR-Cre neurons divided into activated and suppressed subsets of similar size, with or without prior stress. vGAT responses to both sucrose and quinine grew after longer water restriction and scaled with licking, linking them to consummatory engagement rather than taste valence alone. Mice lacking Pdyn in the aIC ate more palatable food even without stress, and KOR ligands changed how mice engage with palatable food. These findings identify an insular–claustral opioid circuit through which stress can shape hedonic feeding.

## 1 INTRODUCTION

Stress and negative affect can increase the consumption of palatable food, even in the absence of hunger (Adam & Epel 2007). Such hedonic feeding, eating for pleasure and reward rather than to meet energy need, can be modeled in rodents: acute stress increases palatable-food intake in mice with no history of food restriction (Anversa et al. 2020). Stress may change not only how much is eaten but also how eating unfolds in time, including how quickly it begins and how often it recurs. Which circuits stress acts on to produce these changes is unclear.

Dynorphin, an endogenous opioid peptide that acts at κ-opioid receptors (KORs), mediates many of the aversive and behavioral consequences of stress (Bruchas et al. 2010). Opioid-receptor antagonism can also reduce palatable-food consumption in people with binge eating, although that work targeted μ-opioid receptors (Cambridge et al. 2013). Where stress-related opioid signals act on the circuits that control eating is not known.

The claustrum is a candidate site. This thin sheet of neurons is reciprocally connected with much of the cortex (Jackson et al. 2020), is engaged by stressors (Niu et al. 2022; Wang et al. 2023) and is enriched in KOR (Bolger et al. 2026). Claustral dynorphin/KOR signaling contributes to depression-like behavior after social stress (Wang et al. 2023), and claustral activity scales with appetitive feeding (de Almeida et al. 2026). Yet the claustrum remains poorly understood, and how its cell types contribute to emotional feeding is unknown. The claustrum contains excitatory projection neurons and local inhibitory interneurons. Inputs or neuromodulators that act differently on these populations could reorganize claustral output without changing its net activity, a change that population measures such as Fos labeling or fiber photometry cannot reveal.

Here we combine behavioral analysis, opioid and glutamate sensors, cell-type-specific one- and two-photon imaging, gene targeting and pose-based behavioral analysis to ask how the claustrum is engaged during stress-induced hedonic feeding. We find that acute stress makes mice start eating sooner and return to food more often, in a manner that involves claustral KOR. Claustral opioid tone falls as eating begins, and Pdyn-expressing neurons of the anterior insular cortex provide an input that recruits claustral neurons. At cellular resolution, inhibitory and KOR-expressing claustral populations are engaged in distinct ways during eating.

## 2 RESULTS

### Stress-induced hedonic feeding involves claustral KOR

To study stress-induced eating without the confound of hunger, we familiarized mice with ad libitum access to chow with a high-palatability diet (HPD; 45 kcal% fat, 30 kcal% sucrose) and tested them in a within-subject design. On each test day, mice either swam for 15 min in 30 °C water (forced swim, FS) or underwent a no-stress control condition (NS), rested for 15 min in the home cage and then had free access to HPD and chow for 1 h (**Fig. 1a**). Mice ate more HPD after FS than after NS, whereas chow intake remained low (**Fig. 1b**).

Stress changed how mice ate as well as how much. After FS, mice approached the HPD almost immediately and returned to it repeatedly throughout the hour (**Fig. 1c**). The latency to the first HPD bout was shorter and the number of feeding bouts was larger (P = 0.002 and P = 0.020, respectively; n = 10 mice; **Fig. 1d,e**), whereas the mean duration of individual bouts did not detectably differ (paired t-test, P = 0.762). Stress therefore acted on when eating started and how often it recurred, rather than on how long each bout lasted. This temporal structure guided our interpretation of the circuit data that follow.

The effect was not specific to one stressor: random footshock also increased HPD intake (**Supplementary Fig. S1I**). Voluntary wheel running did not reproduce the increase (**Supplementary Fig. S1H**), and body weight and 24-h intake did not detectably differ between conditions (**Supplementary Fig. S1B,C**). When chow was the only food available, stress also increased chow intake (**Supplementary Fig. S1A**). The effect is therefore expressed most strongly toward palatable food when a choice is available, but is not exclusive to it.

To find where stress and palatable feeding converge, we examined Fos expression. Fos labeling in the claustrum (CLA) appeared denser after FS+HPD than after NS+HPD, and Fos-positive cells were distributed throughout its rostrocaudal extent (**Fig. 1f,g**). Because the CLA is enriched in KOR, we asked whether claustral KOR contributes to stress-induced feeding. After bilateral injection of AAV-Cre-GFP into the CLA (**Fig. 1h**), KOR-flox mice showed a smaller stress-induced increase in HPD intake than wild-type controls (**Fig. 1i**). Local infusion of the KOR antagonist norBNI into the CLA likewise reduced HPD intake after FS (**Supplementary Fig. S2a**). Together, these results implicate claustral KOR signaling in stress-induced hedonic feeding.

### Opioid dynamics during feeding and an insular Pdyn input to the claustrum

If claustral KOR contributes to stress-induced feeding, opioid tone in the CLA might change as mice eat. To monitor it, we expressed the KOR-based fluorescent sensor kLight (Dong et al. 2024) in CLA KOR-Cre neurons and recorded it with fiber photometry (**Fig. 2a,b**). kLight fluorescence decreased at the onset of HPD feeding bouts (4 mice, 110 bouts; **Fig. 2c**), consistent with a transient fall in local opioid tone as mice began to eat. The decrease was attenuated in sessions after the KOR agonist U50,488 (U50; 5 mg/kg; **Fig. 2b,c**), although these sessions included a partly different set of mice and U50 can also act on the sensor itself. By contrast, a brief tail lift, an acute stressor, increased kLight fluorescence (**Fig. 2d,e**). The claustral opioid signal thus rose with an acute stressor and fell with eating. We next asked where this dynorphin comes from.

We injected a Cre-dependent retrograde AAV into the CLA of Pdyn-Cre mice (**Fig. 2f**). Retrogradely labeled Pdyn neurons were found in the insular cortex, the dorsal endopiriform nucleus, the ectorhinal cortex and the region of the olfactory tract, with more labeled insular cells anteriorly than posteriorly (**Fig. 2g,h**). We focused on the anterior insular cortex (aIC), which integrates gustatory and interoceptive information (Gogolla 2017) and is therefore well placed to convey bodily and emotional state to the CLA.

**Figure 1.**
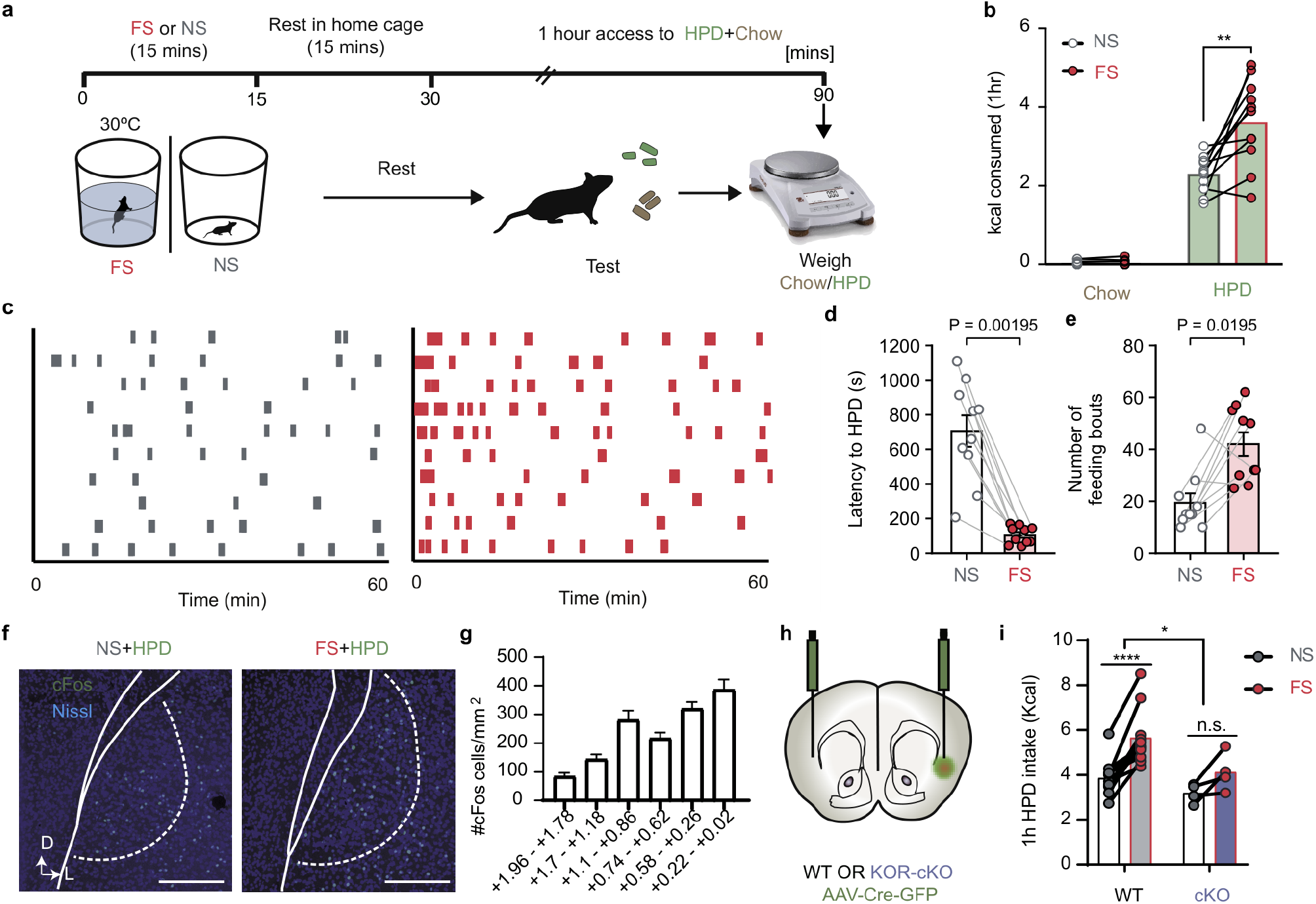
Acute stress reorganizes palatable feeding and engages claustral KOR signaling. **a**, Feeding assay. Ad libitum-fed mice underwent 15 min of forced swim (FS; 30 °C) or the no-stress (NS) condition, rested in the home cage for 15 min, and then received high-palatability diet (HPD) and chow for 1 h. **b**, Energy consumed as chow or HPD after NS and FS. Points represent mice; lines connect repeated measurements from the same animal. **c**, Feeding-bout rasters across the 60-min assay, with one mouse per row. **d**,**e**, Latency to the first HPD feeding bout (**d**) and total annotated feeding bouts (**e**) in the same ten mice under NS and FS. Bars, mean ± s.e.m.; points denote mice and lines connect paired conditions. Two-sided exact Wilcoxon signed-rank tests (all 1,024 sign assignments; average ranks for tied absolute differences): **d**, W = 0, P = 0.001953; **e**, W = 5, P = 0.019531; unadjusted. Panel **b** also includes ten mice. **f**, Representative claustral Fos and Nissl labeling after NS+HPD and FS+HPD. Scale bars, 200 μm. **g**, Fos-positive cell density across rostrocaudal claustral coordinates, reported in cells/mm^2^ (n = 3 mice). This panel shows a spatial profile rather than a stress comparison at each coordinate. **h**, Bilateral claustral AAV-Cre-GFP injections in wild-type (WT) and KOR-flox mice. **i**, One-hour HPD intake after NS and FS in WT and conditional KOR-deletion groups (WT, n = 10; cKO, n = 4); lines connect the same mouse. Gray, NS; red, FS.

**Figure 2.**
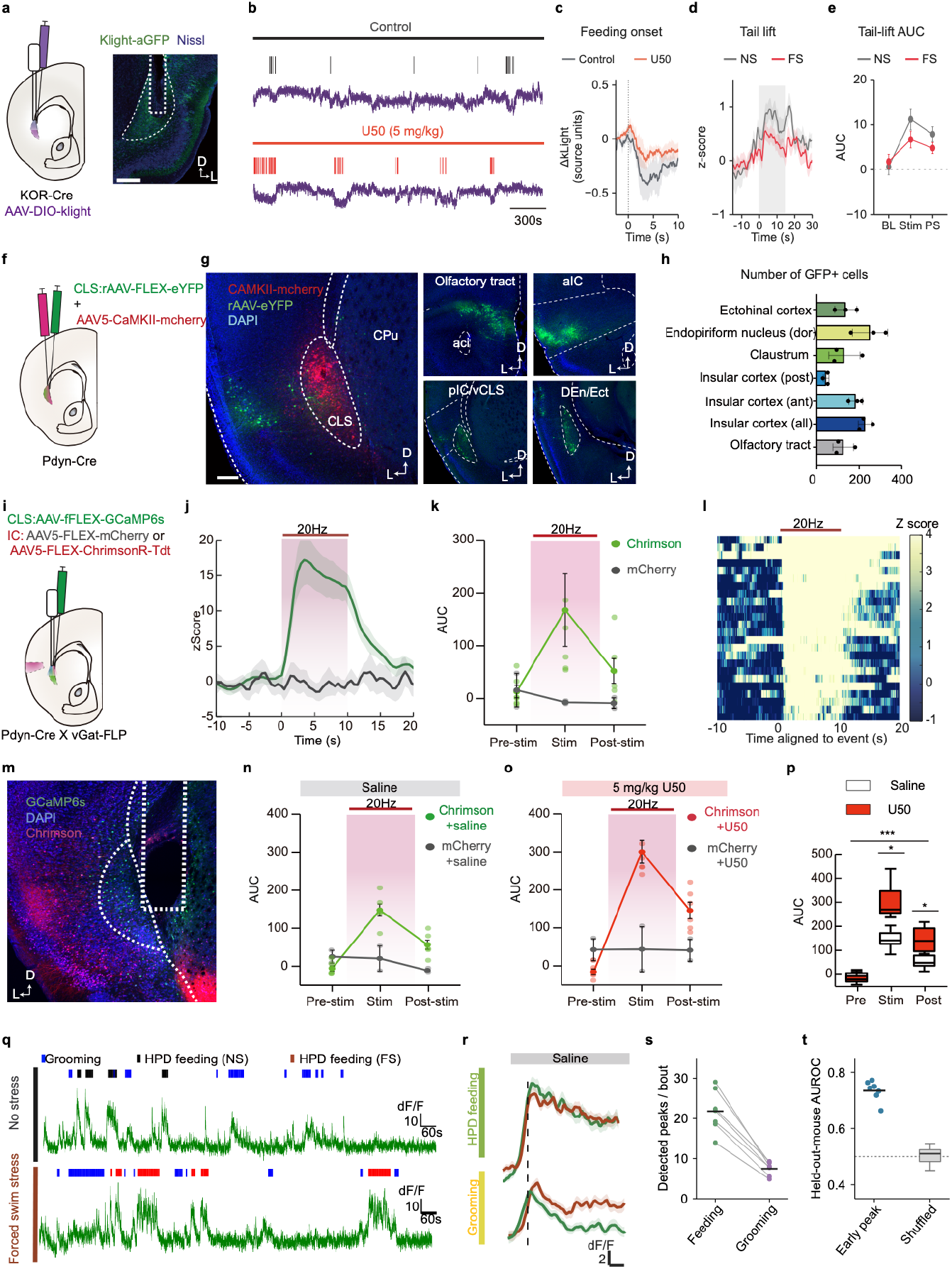
Feeding-associated opioid dynamics and an insular input to claustral vGAT neurons. **a**, kLight targeting in CLA KOR-Cre neurons and representative expression. **b**, Continuous kLight examples with feeding-event ticks in control and U50,488 (U50; 5 mg/kg) sessions; scale bar, 300 s. **c**, Feeding-onset-aligned kLight responses. Each event was baseline-subtracted using −5 to 0 s, averaged within mouse and then across mice; shading indicates s.e.m. across mice. Control: 4 mice, 110 bouts; U50: 6 mice, 233 bouts. The ordinate retains the saved fluorescence source units. **d**,**e**, Tail-lift-aligned kLight responses (**d**) and baseline (BL), stimulation (Stim) and post-stimulation (PS) area under the curve (AUC; **e**), after NS or FS. **f–h**, Retrograde labeling of CLA-projecting Pdyn neurons. Cre-dependent retrograde eYFP and local CaMKII-mCherry were injected into the CLA of Pdyn-Cre mice (**f**). Representative injection-site and projection-source images (**g**) and GFP-positive cell counts across the indicated regions (**h**; n = 3 mice) are shown. **i**,**m**, Cre-dependent ChrimsonR or mCherry in aIC Pdyn neurons and Flp-dependent GCaMP6s in CLA vGAT neurons of Pdyn-Cre × vGAT-Flp mice, with representative expression (**m**). **j–l**, CLA calcium time courses (**j**), pre-/during-/post-stimulation AUC (**k**), and stimulus-aligned heatmap (**l**) during 20-Hz stimulation with 10-ms pulses for 10 s. ChrimsonR, n = 6 mice; mCherry, n = 2. The heatmap color scale and mean-trace ordinate use different display limits. **n–p**, Stimulation-associated AUC after saline (**n**) or U50 (5 mg/kg; **o**), and the corresponding epoch summaries (**p**). **q**,**r**, Continuous vGAT-GCaMP examples and feeding-/grooming-onset summaries after saline under NS and FS. **s**, Fluorescence peaks per bout, summarized within recording and then within mouse: 336 feeding and 623 grooming bouts of at least 10 s from 59 recordings in seven mice. Connected points are paired mouse means; counts depend on bout duration. **t**, Exploratory first-2-s feeding/grooming classification with leave-one-mouse-out evaluation (seven mice; mean AUROC 0.735; within-recording shuffled-label refitting, 1,000 permutations, one-sided P = 0.001).

To test whether this input can drive claustral neurons, we expressed ChrimsonR (or mCherry as a control) in aIC Pdyn neurons and recorded calcium activity in the CLA (**Fig. 2i,m**). In Pdyn-Cre × vGAT-Flp mice, 20-Hz stimulation evoked a large, sustained increase in CLA vGAT population activity in ChrimsonR mice but not in mCherry controls (**Fig. 2j–l**). The same stimulation also recruited CaMKII-expressing claustral neurons (**Supplementary Fig. S3b,c**). vGAT responses were larger after U50 than after saline (**Fig. 2n–p**), suggesting that KOR signaling can modulate this recruitment. aIC Pdyn neurons therefore provide a functional input to both inhibitory and excitatory claustral populations; because optogenetic stimulation engages all transmitters of these neurons, these experiments do not exclude other neuromodulator that might engage this projection. In a real-time place-preference assay, mice expressing ChrimsonR in aIC Pdyn neurons spent more time in the stimulation-paired zone, accompanied by slower movement (**Supplementary Fig. S3d–f**). Stimulation of aIC Pdyn terminals in the CLA produced a similar occupancy bias in a separate cohort and they still prefer the side pared to the light in the subsequent 20-min recording without light (n = 4 mice; **Supplementary Fig. S3g,h**). These observations are consistent with a preference for the stimulation-associated location, indicating a potential positive valence associated with activating aIC Pdyn-CLA projection.

We then asked how claustral populations behave during natural eating. In freely moving mice, CLA vGAT population activity rose at the onset of HPD feeding and remained elevated throughout the bout, after both NS and FS (**Fig. 2q,r**). vGAT activity also rose at the onset of grooming, a self-directed behavior closely linked to stress in rodents (Kalueff et al. 2016), but this response was smaller and decayed quickly (**Fig. 2r**). Feeding bouts contained more calcium peaks than grooming bouts, although this count partly reflects their duration (**Fig. 2s**). In CaMKII-targeted recordings, feeding onset was accompanied by only a small, transient rise in activity, mainly after FS (**Supplementary Fig. S3a**).

Bulk recordings thus revealed robust recruitment of vGAT neurons during feeding but only a small signal from CaMKII-expressing neurons. A small bulk signal could mean that excitatory neurons are weakly engaged, or that individual neurons respond strongly but in opposite directions, cancelling in the average. And because vGAT neurons also responded during grooming, bulk activity could not tell us which aspects of feeding they encode. To address both questions, we turned to single-cell imaging.

### vGAT and KOR-Cre neurons respond to feeding in distinct ways

We imaged single CLA neurons with a head-mounted miniscope in freely moving mice, targeting two genetically defined populations: vGAT neurons (vGAT-Flp mice expressing Flp-dependent GCaMP6s; **Fig. 3a**) and KOR-Cre neurons, the cells on which dynorphin can act directly (KOR-Cre mice expressing Cre-dependent GCaMP6s; **Supplementary Fig. S4a**). Each mouse was recorded in the feeding assay after NS and after FS (three mice per population).

**Figure 3.**
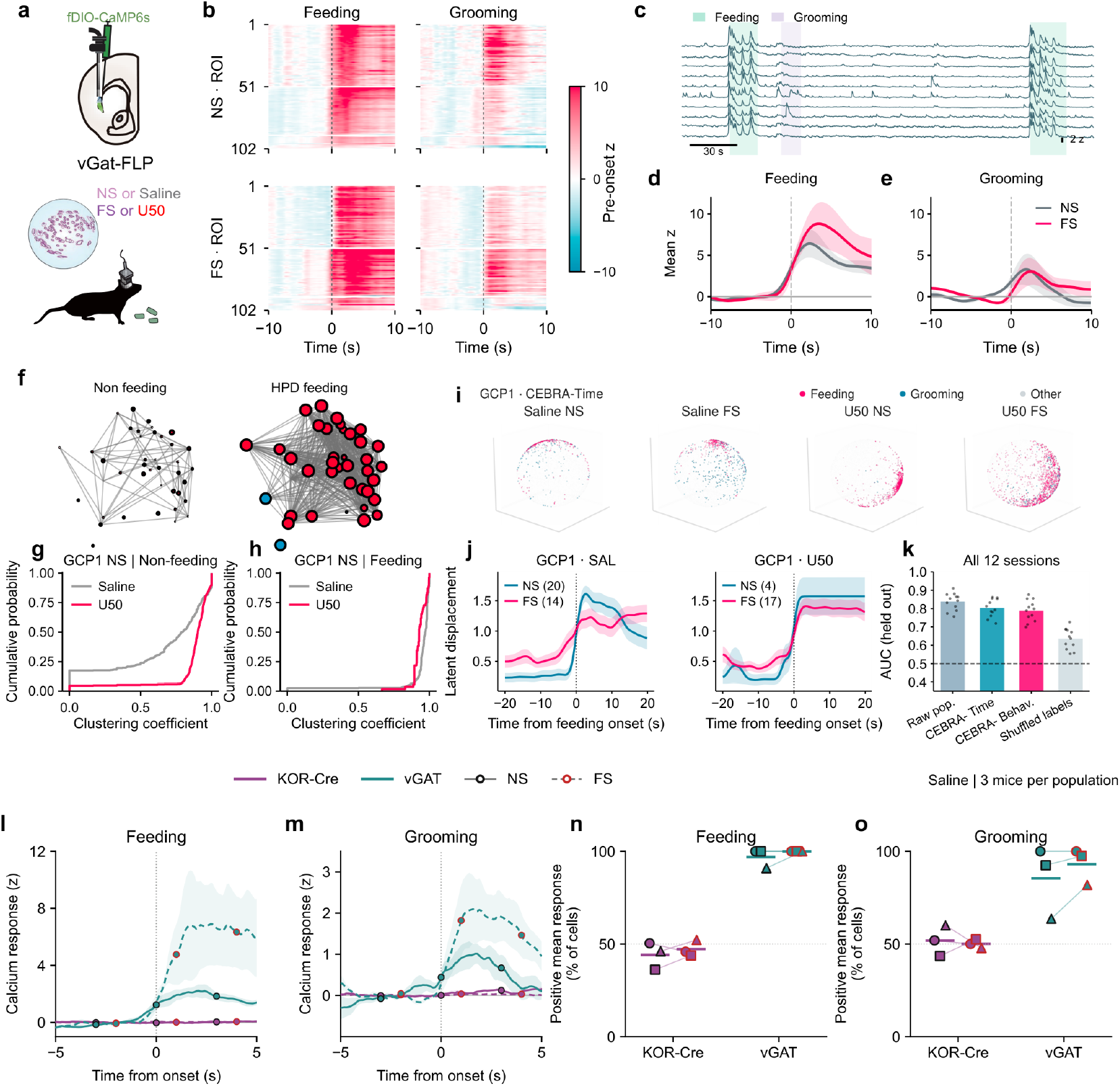
Distinct claustral population responses during feeding and grooming. a–e, Freely moving one-photon imaging of CLA vGAT neurons expressing Flp-dependent GCaMP6s (**a**). Behavior-onset heatmaps retain 102 displayed ROI rows under saline NS and FS (**b**). **c**, Example calcium traces with feeding and grooming epochs; scale bars, 30 s and 2 z. **d**,**e**, Feeding- and grooming-onset means ± SEM across three mouse means, using the original processing. Gray, NS; magenta, FS. **f–h**, Representative nonfeeding/feeding coactivity graphs from GCP2 under saline NS (**f**), and GCP1 NS clustering-coefficient distributions for saline versus U50 during nonfeeding (**g**) or feeding (**h**). Graph edges denote positive Pearson correlations above 0.5. These are single-animal descriptions of functional coactivity. **i–k**, GCP1 CEBRA-Time representations for saline NS/FS and U50 NS/FS (**i**; 3,500 displayed points per session). Each session was fitted independently. **j**, Feeding-aligned trajectories from behavior-supervised embeddings; displayed bout numbers are saline NS/FS, 20/14, and U50 NS/FS, 4/17. Bands summarize bouts within GCP1. **k**, Exploratory decoding summaries across 12 sessions from three mice; points denote session summaries. The original artwork and its labels are retained. **l–o**, Common downstream comparison of KOR-Cre and vGAT populations under saline (n = 3 mice per population). **l**,**m**, Equal-mouse mean ± between-mouse SEM during feeding (**l**) and grooming (**m**). Each event is baseline-subtracted over −5 ≤ t < 0 s; responses are scaled by each cell’s pooled within-event baseline sample SD. Cells are averaged within recording, recordings within mouse, then mice equally. Feeding and grooming retain separate y-axis ranges shared across populations and states. Purple, KOR-Cre; teal, vGAT. NS uses solid lines/black outlines; FS uses dashed lines/red outlines. Sparse trace markers identify state, not independent samples. **n**,**o**, Cells with a positive mean response over 0 ≤ t ≤ 5 s after feeding (**n**) or grooming (**o**) onset, summarized per mouse. Symbols denote mice, lines pair NS/FS within mouse, and short bars show equal-mouse means.

Eating recruited CLA vGAT neurons almost uniformly. Nearly all vGAT neurons increased their activity at feeding onset, after both NS and FS (**Fig. 3b–d**). The same neurons also responded at grooming on-set, but less strongly (**Fig. 3b,e**).

KOR-Cre neurons behaved differently. At feeding onset, some increased and others decreased their activity, so that the population average was much smaller (**Fig. 3l**; **Supplementary Fig. S4b**). To compare the two populations directly, we reprocessed both datasets with identical event windows, normalization and inclusion criteria, weighting each mouse equally (**Fig. 3l–o**). After NS, 97% of vGAT neurons but only 44% of KOR-Cre neurons had a positive mean response in the 5 s after feeding onset; after FS, the values were 100% and 47% (**Fig. 3n**). Grooming responses were smaller and more variable, but still differed between the populations (**Fig. 3m,o**). Eating therefore recruits vGAT neurons as a near-uniform block but divides KOR-Cre neurons into activated and suppressed subsets of similar size. This organization was present with and without prior stress, and population averages, including bulk photometry, would conceal it.

Feeding also changed coordinated activity among vGAT neurons. In example mice, coactivity graphs were denser during feeding than outside it, and U50 increased clustering outside feeding toward the level seen during feeding (**Fig. 3f–h**). Low-dimensional embeddings of population activity (CEBRA; Schneider et al. 2023) showed an abrupt shift in population state at feeding onset under both saline and U50 (**Fig. 3i,j**), and exploratory decoding analyses are summarized in **Fig. 3k**. These single-animal analyses are descriptive, but they illustrate that feeding engages vGAT neurons as a coordinated population.

KOR activation altered KOR-Cre responses without changing their overall balance. After systemic U50, the fractions of KOR-Cre neurons classified as activated or suppressed at feeding onset were similar to those in control sessions (**Supplementary Fig. S4c**), but neurons that had responded in control sessions largely lost their responses (**Supplementary Fig. S4d**). Opioid signaling may therefore change which KOR-Cre neurons respond during eating rather than how many; we return to this possibility in the Discussion.

### Claustral vGAT responses to palatable and aversive tastes scale with internal state and licking

What aspect of feeding do vGAT neurons signal? Their recruitment during free feeding could reflect the valence of the food, its sensory properties, the animal’s internal state, such as stress or motivation, or the act of consumption itself, factors that are entangled when mice eat freely. To separate them, we used two-photon imaging in head-fixed mice, which allowed us to deliver tastes of opposite valence, follow the same neurons as internal state changed and measure consumption lick by lick (**Fig. 4a**). Because head-fixed mice cannot eat freely, we manipulated internal state with water restriction: mice received sucrose and quinine after overnight water restriction (day 1) and again after two days of water restriction (day 3). Mice licked more for both tastants on day 3 (**Fig. 4a**, inset). We tracked the same 124 neurons across both days and both tastants in three mice (**Fig. 4b**).

**Figure 4.**
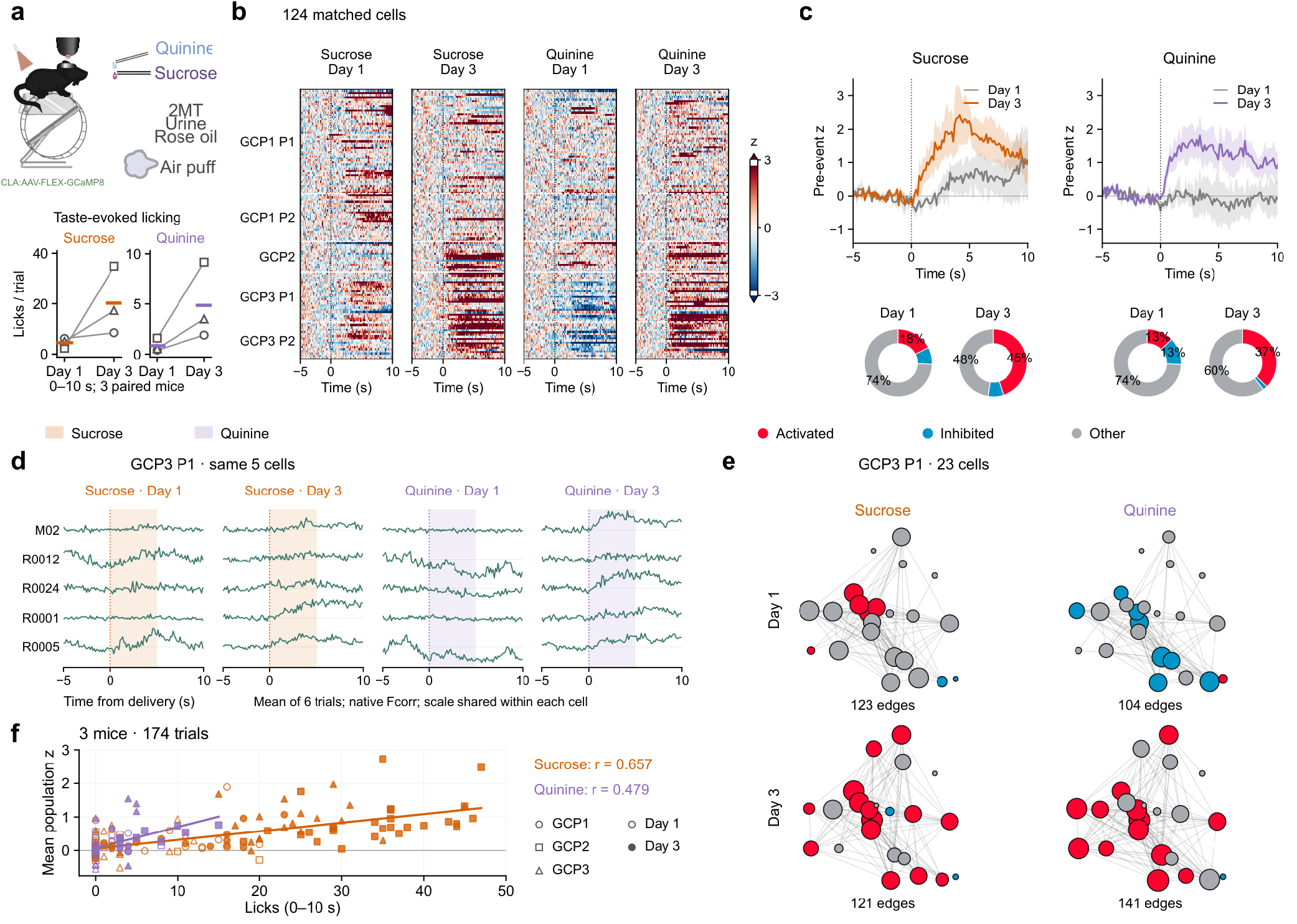
Longitudinal claustral vGAT responses during sucrose and quinine consumption. **a**, Head-fixed two-photon imaging with taste delivery and a wheel. The behavioral inset shows each mouse’s mean number of licks during 0–10 s after taste delivery on Day 1 and Day 3; lines connect mice, and colored bars show the equal-mouse mean (three mice; 191 complete behavior trials, including zero-lick trials). Circles, squares and triangles identify GCP1, GCP2 and GCP3. Taste-specific y-axis scales differ. The imaging dates were separated by one intervening day of water restriction. **b**,**c**, Taste-aligned responses of 124 matched ROIs from three mice (GCP1, 70; GCP2, 14; GCP3, 40), with identical cell order across heatmaps (**b**), equal-mouse mean ± s.e.m. traces and mouse-averaged response-class proportions (**c**). Fluorescence was neuropil-corrected as F − 0.7Fneu, averaged across selected trials within cell, and normalized to the mean and standard deviation of the event-averaged −5 to 0 s baseline. The primary set contained 10, 6 and 6 trials per taste and day for GCP1–3, respectively, with complete windows shared across recording planes. Heatmap limits are z = ±3. Activated, inhibited and other denote mean 0–10-s z > 1, < −1 and between those thresholds, respectively; these are descriptive amplitude classes, not significance tests. **d**, Baseline-subtracted, six-trial-average native fluorescence responses from the same five GCP3 plane-1 cells. One display gain per cell is held constant across all four conditions; amplitudes are not comparable across cells. Shaded blocks mark the first 5 s after delivery. **e**, Coactivity graphs of the same 23 GCP3 plane-1 cells, using OASIS-reconstructed calcium from four fixed taste epochs totaling approximately 80.34 s per condition. Edges indicate positive Pearson r > 0.2; positions are anatomical, node diameter increases with degree, and colors denote the response classes in **c**. These are one-mouse descriptive coactivity examples, not anatomical connectivity. **f**, Lick count versus mean population calcium response over 0–10 s, for 174 calcium-QC-valid physical trials (126 sucrose, 48 quinine) from three mice. Each cell/trial was normalized to its own −5 to 0 s baseline before averaging across reviewed cells. Open/filled markers denote Day 1/Day 3; shapes identify mice.

On day 1, most neurons responded weakly: sucrose activated 18% of neurons, and quinine activated 13% and inhibited 13% (**Fig. 4b,c**). On day 3, the same neurons responded more strongly to both tastants: 45% were activated by sucrose and 37% by quinine, and few were inhibited by either (**Fig. 4b,c**). Quinine responses thus shifted from mixed to predominantly positive, as illustrated by example neurons tracked across all four conditions (**Fig. 4d**). In an example mouse, coactivity among the same 23 neurons also differed across tastes and days (**Fig. 4e**).

Because licking also increased on day 3, we asked whether neural activity tracked consumption itself. Across 174 trials from three mice, trial-by-trial population activity scaled with the number of licks for both sucrose (r = 0.66) and quinine (r = 0.48), and the association remained after removing differences between mice and days (r = 0.45 and 0.25; **Fig. 4f**). vGAT activity therefore relates to how much the animal consumes, for a palatable and an aversive tastant alike, rather than to taste valence alone.

In the same preparation, vGAT neurons also responded to non-gustatory stimuli, including an air puff, infrared stimulation and several odors (**Supplementary Fig. S5**). Together with the freely moving recordings, these results suggest that claustral vGAT neurons respond broadly across sensory modalities and are recruited during consumption in proportion to internal state and consummatory engagement.

### Peptide and glutamate components of the insular input

aIC Pdyn neurons could influence the CLA through dynorphin, glutamate or both. CLA-projecting aIC Pdyn neurons expressed vGluT1 (**Fig. 5a**), so this input is both peptidergic and glutamatergic. Consistent with dual signaling, simultaneous recording of kLight and the glutamate sensor iGluSnFR (Marvin et al. 2013) in an independent vGAT-targeted cohort showed that, at feeding onset, kLight fluorescence decreased while iGluSnFR fluorescence increased (four mice; **Fig. 5b–e**). Eating thus coincides with a fall in local opioid tone and a rise in glutamatergic signaling in the CLA, and the kLight decrease replicated that seen in the KOR-Cre cohort (**Fig. 2c**). These local signals cannot by themselves be attributed exclusively to the aIC input.

**Figure 5.**
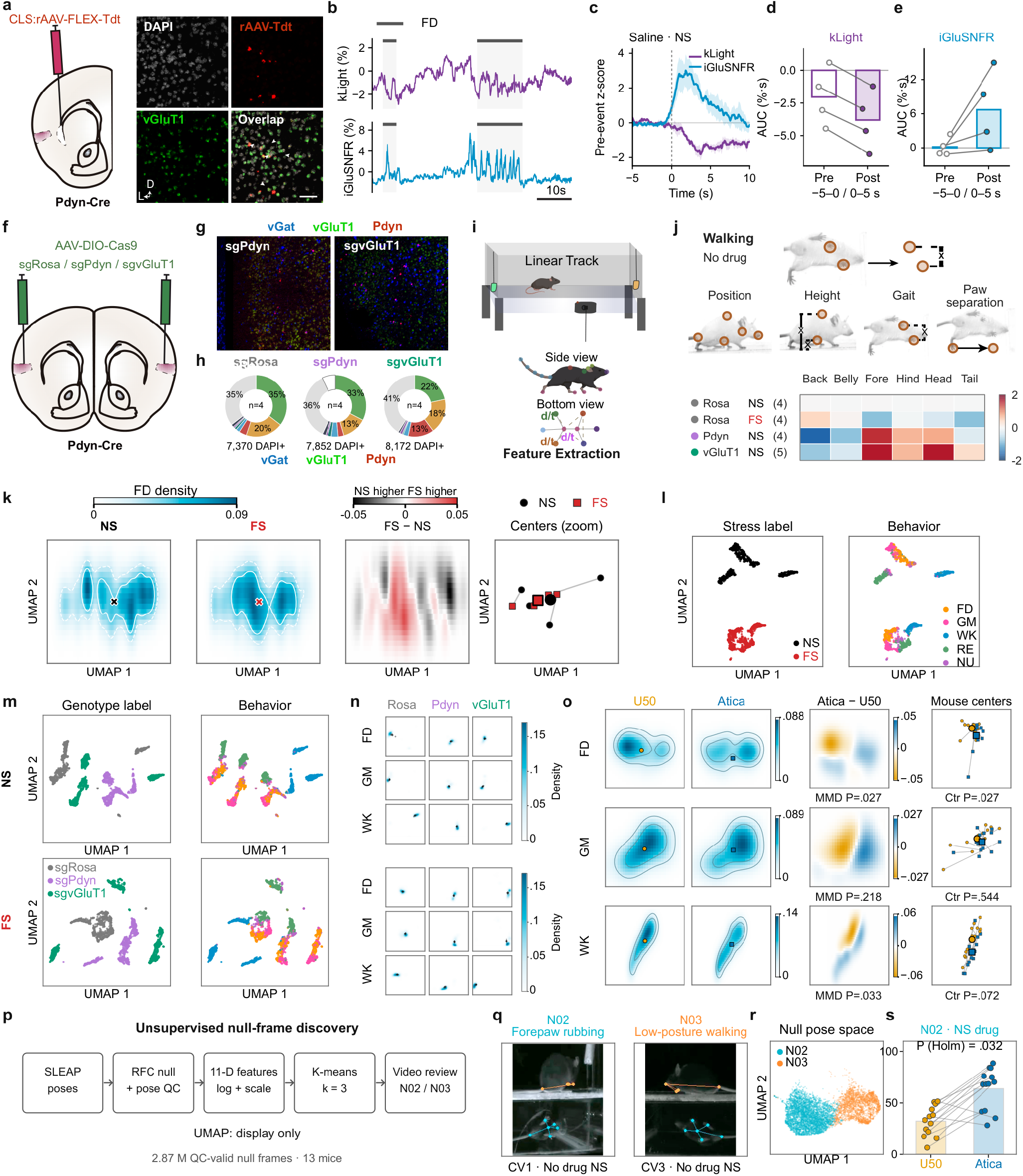
Transmitter-targeted perturbations and pose-based phenotyping of the insular–claustral circuit. **a**, Retrograde labeling of CLA-projecting Pdyn neurons and representative reporter/vGluT1 labeling. **b–e**, Simultaneous claustral kLight and iGluS-nFR recordings in the separate vGAT-targeted sensor cohort under saline and NS. **b**, Continuous example with feeding epochs; scale bar, 10 s. **c**, Event-aligned responses normalized to each event’s −5 to 0 s baseline, averaged within mouse and then across four mice (113 feeding onsets); shading, s.e.m. across mice. **d**,**e**, Native-fluorescence AUC in −5 to 0 s and 0 to 5 s for kLight (**d**) and iGluSnFR (**e**), in percentage-point∙s. Bars are mouse means; connected points are paired mouse measurements. No inferential tests are shown for **d/e. f–h**, Cre-dependent Cas9 and guide RNAs targeting Rosa26 (control), Pdyn or Slc17a7/vGluT1 in aIC Pdyn neurons (**f**), representative multiplexed RNA images (**g**), and marker-defined composition (**h**). Donuts average four mouse-specific composition vectors per group; total DAPI-positive object counts are 7,370, 7,852 and 8,172 for sgRosa, sgPdyn and sgvGluT1. Colored sectors denote mutually exclusive combinations of vGAT, vGluT1 and Pdyn labeling; hatched sectors indicate unavailable vGAT classification. These provisional marker calls characterize composition and do not establish editing efficacy. **i**,**j**, Two-view linear-track tracking and feature extraction (**i**), with no-drug walking features (**j**) standardized to the four sgRosa-NS mouse means. Group sizes are sgRosa NS/FS, four paired mice; sgPdyn NS, four; sgvGluT1 NS, five. Height, left–right paw separation and lateral head/tail deviation are normalized pose measures; paw separation is not stride length. **k**, Conditional feeding densities and paired centers for four sgRosa mice after NS and FS in a fixed behavior-supervised map. **l**, Stress-supervised embedding and behavioral recoloring. **m**,**n**, Separate genotype-supervised NS and FS embeddings (**m**) and conditional feeding, grooming and walking densities (**n**), with 4/4/5 mice in the three guide groups. These label-informed displays are descriptive; distances between separately fitted maps are not comparable. **o**, NS U50 versus aticaprant comparisons in 13 paired mice. Densities are normalized within mouse and condition. Inference used the original 17-dimensional features, with all 8,192 whole-mouse paired-label permutations and Holm correction across five behaviors × two endpoints. Adjusted P values for distribution (MMD)/centroid comparisons were 0.027/0.027 for feeding, 0.218/0.544 for grooming and 0.033/0.072 for walking. **p–r**, Unsupervised partitioning of 2.87 million QC-valid frames left unassigned by the behavior classifier, using 11 pose features and k-means (k = 3); UMAP is for display only. Example frames illustrate candidate N02 forepaw rubbing and N03 low-posture walking. **s**, N02 occupancy among all QC-valid unassigned frames, including N01, after NS U50 or aticaprant (13 paired mice; mean bars and connected mouse points). Holm-adjusted P = 0.032 from paired mouse-label permutations, corrected over 24 endpoint/contrast tests. The N02 association weakened to P = 0.056 with stricter pose-score QC.

To ask whether insular dynorphin constrains feeding, we deleted Pdyn from the aIC in an independent cohort, injecting AAV-Cre bilaterally into the aIC of conditional Pdyn-knockout mice (Pdyn-cKO), with wild-type mice as controls. Without stress, Pdyn-cKO mice ate nearly twice as much HPD as controls (0.78 ± 0.04 versus 0.41 ± 0.07 g; n = 5 and 6 mice; P = 0.004; **Supplementary Fig. S2d**). Stress still increased intake in every cKO mouse, so loss of insular Pdyn did not abolish the response to stress. These data suggest that insular dynorphin restrains palatable-food intake in the absence of stress, although the two groups also differed in age, which needs to be addressed in a matched cohort.

To separate the two transmitters within the same neurons, we used CRISPR–Cas9 to target Pdyn or vGluT1 (Slc17a7) selectively in aIC Pdyn neurons, injecting Cre-dependent Cas9 into the aIC of Pdyn-Cre mice together with guide RNAs against Rosa26 (sgRosa, control), Pdyn (sgPdyn) or vGluT1 (sgvGluT1) (**Fig. 5f**). Multiplexed in situ hybridization characterized the cellular composition of each guide group (**Fig. 5g,h**) but has not yet established loss of the targeted transcripts. In this exploratory cohort, the stress-induced increase in intake tended to be smaller in sgPdyn mice, consistent with the cKO result, whereas sgvG-luT1 mice still showed an increase; the sgPdyn pattern depended on whether one questioned low-intake record was included (**Supplementary Fig. S2b,c,e,f**).

Because stress changed how mice ate rather than only how much (**Fig. 1c–e**), we also developed a pose-based approach to quantify the structure of behavior. Mice were filmed from side and bottom views on a linear track, body parts were tracked with SLEAP (Pereira et al. 2022) and posture and gait features were extracted for each labeled behavior (**Fig. 5i,j**). In control mice, stress shifted the distribution of feeding postures (**Fig. 5k**), and embeddings informed by stress or guide labels separated the conditions (**Fig. 5l–n**). Because these labels contribute to the fitted maps, we tested condition effects in the original feature space, where guide-group differences did not survive correction for multiple comparisons.

KOR ligands, by contrast, changed how mice ate. In 13 mice tested after U50 and after the KOR antagonist aticaprant, feeding postures differed between drugs in both distribution and center (Holm-adjusted P = 0.027), as did walking postures (P = 0.033), whereas grooming did not (**Fig. 5o**). Unsupervised clustering of 2.87 million quality-controlled frames left unlabeled by the behavior classifier revealed candidate behaviors, including forepaw rubbing and low-posture walking (**Fig. 5p–r**). Forepaw rubbing was more frequent after aticaprant than after U50 (Holm-adjusted P = 0.032; **Fig. 5s**), an association that weakened under stricter tracking quality control. These measurements make it possible to relate peptide signaling to the fine structure of feeding, beyond total intake.

## 3 DISCUSSION

We show that a single acute stressor makes mice without food restriction start eating palatable food sooner and return to it more often, and that claustral KOR contributes to this effect. Within the claustrum, opioid tone falls as eating begins, and Pdyn-expressing neurons of the anterior insula provide an input that recruits claustral neurons. At cellular resolution, two genetically defined claustral populations are engaged in distinct ways: vGAT neurons are recruited almost uniformly, whereas KOR-Cre neurons split into activated and suppressed subsets of similar size. The behavioral effect of stress was observed in both sexes, with a more bimodal distribution of intake in females.

Fos labeling and photometry indicate that the claustrum is activated during stress-induced feeding, but cellular imaging shows that this net activation contains two distinct modes of engagement. vGAT neurons are recruited during feeding, are driven by insular Pdyn input and respond to both sucrose and quinine in proportion to consumption. They may therefore signal engagement in a consummatory act, and its intensity as scaled by internal state, rather than the identity or valence of a stimulus. This extends recent evidence that claustral activity scales with appetitive feeding (de Almeida et al. 2026) by resolving how genetically defined populations contribute. The balanced split of KOR-Cre neurons, by contrast, means that eating changes which of these neurons are active rather than simply how active the population is. One possibility is that broadly recruited inhibitory neurons, together with direct insular excitation, sculpt claustral output into opposing subsets.

Opioid signaling is positioned to act on both sides of this circuit. KOR expression is enriched in excitatory claustral neurons (Bolger et al. 2026). Under U50, KOR-Cre neurons kept the same balance of activated and suppressed cells, but neurons that had responded in control sessions largely stopped responding, and vGAT responses to insular input became larger. Dynorphin may therefore reshape which KOR-Cre neurons participate in eating while leaving their overall balance intact. Distinguishing this from a global change in gain or regression to the mean will require comparing ensemble membership across sessions, both against chance and against repeated control recordings.

The temporal structure of stress-induced feeding suggests a working model. Eating lowered claustral opioid tone in two independent cohorts (**Fig. 2c** and **Fig. 5b–e**), whereas an acute stressor raised it (**Fig. 2d,e**). We propose that the fall in opioid tone during eating acts as a local feedback signal of consumption. If stress elevates claustral dynorphin tone or blunts its fall, each bout would deliver less of this signal, and more bouts would be needed to reach the same state, matching the earlier and more frequent eating we observed (**Fig. 1c–e**). Removing dynorphin from the insular input would weaken the signal itself, so that unstressed mice would eat more, as Pdyn-cKO mice did. The model predicts that the feeding-related kLight decrease should be smaller after stress and that the timing of eating should depend on the size of this decrease. Our current recordings do not yet resolve the first prediction: within-mouse comparisons of NS and FS sessions were mixed, with a shallower decrease after stress in two of four mice and a deeper one in the other two. Because mean bout duration did not change, the model concerns the initiation and recurrence of eating rather than when a bout ends.

The loss-of-function results also do not yet align into a single ligand–receptor mechanism. KOR deletion or blockade in the CLA attenuated stress-induced eating, whereas Pdyn deletion in the aIC increased intake without stress. These interventions differ in anatomy, cell targeting and validation, and aIC Pdyn neurons project beyond the CLA, while other regions, notably the endopiriform nucleus, also supply Pdyn input to it. Manipulating ligand and receptor within the same pathway will be needed to reconcile them.

Two approaches developed here may be useful beyond this circuit. CRISPR targeting within a molecularly defined projection population offers a way to separate a co-released peptide from a fast transmitter in vivo, a persistent problem for peptidergic circuits. Pose-based feature extraction quantifies how animals eat, not only how much: KOR ligands changed feeding posture in a way that total intake would not reveal, and unsupervised analysis of unlabeled frames identified candidate behaviors that standard classifiers miss.

Several limitations qualify these conclusions. KOR agonists have sex-dependent effects (Abraham et al. 2018); whether the population organization differs in females is unknown. Response fractions describe the sign of each neuron’s mean response rather than statistically significant activation, and KOR-Cre neurons are not a purely excitatory population. kLight reports opioid tone indirectly, and U50 can act on the sensor itself. In the two-photon experiments, water restriction, experience and licking changed together between days. Optogenetic stimulation engages all transmitters released by aIC Pdyn neurons, and the place-preference result is accompanied by slower movement. The Pdyn-cKO and control groups differed in age, the CRISPR experiments remain exploratory until loss of the targeted transcripts is confirmed, and drug order was not fully balanced in the pose experiments.

Together, our findings place the claustrum, a poorly understood structure, within the circuitry that links emotional state to eating. A stressor and a meal move claustral opioid tone in opposite directions. An input from Pdyn- and vGluT1-expressing neurons of the anterior insula recruits both inhibitory and excitatory claustral neurons, and stimulation was associated with preference for the stimulation-paired location. Within the claustrum, inhibitory and KOR-expressing populations are both engaged during eating but are organized in distinct ways, a distinction invisible to bulk measures of activity. By combining neuromodulatory and fast transmission, a single cortical input may thus both drive and tune claustral output. Stress-related eating is common, and opioidreceptor antagonists can reduce palatable-food consumption in people with binge eating (Cambridge et al. 2013). Defining how peptide and glutamate signals from the insula act on distinct claustral populations should help explain how stress reshapes not only how much we eat, but when and how we eat.

## 4 METHODS

### Animals

Adult wild-type C57BL/6J (Jackson Laboratory #000664), KOR-Cre (Jackson Laboratory #035045), *Pdyn*-IRES-Cre (Jackson Laboratory #027958), *KOR*^*fl/fl*^ (Jackson Laboratory # 037512), vGAT-IRES-Cre (Slc32a1tm2(cre)Lowl/J; Jackson Laboratory #028862) are used in this study. Mice were group housed, given access to food and water ad libitum, and maintained on a 12:12 hr reverse light:dark cycle (lights off at 9:00 AM and on at 9:00 PM). All animals were kept in an isolated and sound-attenuated holding facility within the lab and adjacent to behavior rooms one week prior to surgery, post-surgery, and throughout the duration of the behavioral assays to minimize stress. Males and female mice were used in all experiments. All procedures were approved by the Animal Care and Use Committee of Washington University and the University of Washington and conformed to US National Institutes of Health guidelines.

### Binge Eating Paradigm

#### Stress, no stress, and empty bucket experiments

Mice were individually housed or housed with a partner in a divided cage containing a custommade acrylic divider with hole perforations (American Products, Ltd., Auburn, WA and Curbell Plastics, Inc., Arlington, TX). Isolating each mouse in this way allowed accurate food intake measurements in the home cage. Mice were transferred to single-housing or double housing with the cage divider at least 1 week prior to commencing experiments. Two days before forced swim stress, body weight measurements were taken, and mice were given pre-weighed standard chow (PicoLab Rodent Diet 20^®^, #5053, St. Louis, MO) and custom-made high fat, high sugar food (termed “highly palatable diet” or HPD; 45 kcal% fat, 30 kcal% sucrose, 4.7 kcal/g; Research Diets, Inc., New Brunswick, NJ, ID: D08112601). The pre-weighed diets were given in ample supply to allow choice *ad libitum* feeding for 24 hours.

After 24 hours of free choice feeding, chow and HPD were weighed separately and recorded. Afterwards, only chow was placed back in the home cage. The next day, mice were randomly assigned to stress, no stress or empty bucket groups. In all conditions, prior to feeding assay, all food was removed from the home cage. In the no stress groups, pre-weighed chow, or HPD+chow were placed in the cage and mice were returned to their usual space on the cage rack for 1 hour. In the stress and empty bucket groups, mice were brought to a separate testing room and underwent forced swim stress (mice placed in a 30ºC, 1 L clear plastic bucket of water filled 3/4 of the way for 15 minutes) or foot shock stress (5 minutes of no shocks in chamber, followed by 0.8 mA shocks of 0.5 seconds, randomly administered over a 15 minute period with a 40 second variable interval (10-70 sec range); mice received 20-22 shocks in total) (Redila and Chavkin, 2008, Psychopharmacology) or were placed in an empty 1-L clear plastic bucket. For forced swim sessions, an overhead camera was mounted above water buckets to record immobility time, which was later analyzed with EthoVision software.

After 15 mins of stress, mice who underwent forced swim were patted dry with a paper towel and placed in a clean cage with absorbent paper towels for 5 minutes, and then returned to their home cage for 10 additional minutes. Mice who underwent foot shock were simply placed back in their home cage for 15 minutes. After the 15 minute “rest” period following stress, pre-weighed chow or HPD+chow were placed in the home cage of the stressed mice and mice were returned to their usual space on the housing room rack for 1 hour. Video recordings of feeding behavior were taken during the 1 hour testing period, and were later analyzed and scored by an observer for feeding behavior. The onset of a feeding bout was defined by the first bite of food, and the offset of a feeding bout was defined by cessation of chewing after the last bite taken. After an hour, food weight was measured and recorded.

#### Wheel-running experiments

Mice were individually housed and given access to a small running wheel (Amazon, Ware Manufacturing, ASIN: B06XH5RFFR) placed in the middle of the cage. Mice were left alone for 1 week to allow acclimatization and use of the running wheel. After 1 week access, the running wheels were removed from the home cage, and mice were given ample pre-weighed Chow and HPD for 24 hours for choice *ad libitum* feeding. After 24 hours, food was removed and weighed. Only chow was placed back in the home cage. The following day, mice were brought to a separate testing room, and a running wheel was placed in the home cage for 1 hour. Overhead cameras were mounted to record running activity, which was later watched by an observer and hand-scored for time spent actively running on the wheel. “Runners” were classified as running at least 15 minutes total of the hour tested, and “Stationary” participants were counted if they ran <15 mins during the hour long test period.

### Stereotaxic Surgery

After the animals were acclimated to the holding facility for at least seven days, the mice were anesthetized in an induction chamber (2% isoflurane) and placed into a stereotaxic frame (Kopf Instruments, Model 1900) where they were maintained at 1-2% isoflurane. Mice were then injected using a Nanoject II (Drummond Scientific). Mice were injected in the Calustrum (AP +0.7 mm, ML +2.75 mm, DV – 4.0 mm relative to bregma) with 150-400nL virus at a rate of 100 nL/min. Injections were unilateral for tracing and fiber photometry experiments; unilateral for optogenetic or bilateral for chemogenetic experiments. The needle was slowly removed from the brain 10 min after cessation of injection to allow for diffusion. For fiber photometry and optogenetic experiments, mice were also implanted with a 400-µm fiber optic (Doric Inc., MFC_400/430-0.48_MF2.5_FLT) or custom-made optical fibers of ≥70% optical efficiency in the same surgery. For mice in the aIC photostimulation study to examine effect on the CLA, in addition to the CLA injections as before, 500nl of the virus expressing ChrimsonR were injected using coordinate (AP +0.8 mm, ML +3.27 mm, DV – 4.0 mm relative to bregma) to target aIC Mice received carprofen and recovered for at least 6 weeks prior to behavioral testing, permitting optimal expression of the virus. For fiber photometry experiments, fiber optics were placed unilaterally. Fiber optic implants were secured using Metabond (Parkell #S380).

### Viruses

AAV-DJ-EF1α-DIO-GCaMP6f Stanford Gene Vector and Viral Core

AAV5-EF1α-DIO-mcherry UNC Vector Core

AAV5/Syn-FLEX-ChrimsonR-TdTomato UNC Vector Core

AAV5-CAG-DIO-kLight1.3a Lin Tian (Tian et al. 2023)

AAV-DJ-EF1α-DIO-GCAMP6s Stanford Gene Vector and Viral Core

### Histology

Briefly, mice were euthanized with sodium pentobarbital and transcranial perfused with 4% paraformaldehyde (PFA), post-fixed for 1-3 days in 4% PFA, and then cryo-protected in 30% sucrose. Brain sections (30-100µm) were collected and kept in 0.1M phosphate buffer at 4 until mounting. Sections were mounted with VectaShield Vibrance Hardset mounting medium (Vector Laboratories) with DAPI, and coverslips placed. For enhancing klight expression, tissues are sliced using Vibratome for 100um thickness. Free-floating sections were washed in 0.1M PBS for 3 × 10 minutes intervals. Sections were then placed in blocking buffer (0.5% Triton X-100 and 3% BSA in 0.1 M PBS) for 1 hr at room temperature. Sections were incubated with primary antibody (aGFP Sigma 1:1000) at 4°C overnight. After 3 × 10 minute 0.1M PBS, sections were incubated in Alexa Fluor G-R 488 secondary antibodies (1:1000) and NeuroTrace (1:400, 435/455 blue fluorescent Nissl stain, Invitrogen #N21479) for 1 hour, followed by 3 × 10 minute 0.1M PBS then 3 × 10 minute 0.1M PB washes. Sections were then mounted and coverslipped. All histology Images were acquired on a Confocal microscope (Olympus Fluoview 3000) for 10um step size with either a 10X or 20X objective.

### Electrophysiology

Coronal brain slices were prepared at 250 μM on a vibrating Leica VT1000S microtome using standard procedures. Mice were anesthetized with Euthasol (Virbac, Westlake Texas), and transcardially perfused with ice-cold and oxygenated cutting solution consisting of (in mM): 93 N-Methyl-D-glucamine (NMDG), 2.5 KCL, 20 HEPES, 10 MgSO4 7H20, 1.2 NaH2PO4, 0.5 CaCl2 2H20, 25 glucose, 3 Na^+^-pyruvate, 5 Na^+^-ascorbate, and 5 N-acetylcysteine. Following collection of coronal sections, the brain slices were transferred to a 34°C chamber containing oxygenated cutting solution for a 10-minute recovery period. Slices were then transferred to a holding chamber consisting of (in mM) 92 NaCl, 2.5 KCl, 20 HEPES, 2 MgSO47H20, 1.2 NaH2PO4, 30NaHCO3, 2 CaCl22H20, 25 glucose, 3 Na-pyruvate, 5 Na-ascorbate, 5 N-acetylcysteine and were allowed to recover for ≥ 30 min. For recording, slices were perfused with oxygenated artificial cerebrospinal fluid (ACSF; 31-33°C) consisting of (in mM): 113 NaCl, 2.5 KCl, 1.2 MgSO4-7H20, 2.5 CaCl2-6H20, 1 NaH2PO4, 26 NaHCO3, 20 glucose, 3 Na^+^-pyruvate, 1 Na^+^-ascorbate, at a flow rate of 2-3ml/min. Stocks of 5mM U69,693 were made in ddH_2_0 and added to either the HEPES or ACSF in a 1:5000 ratio for a final concentration of 1μM,.

Claustrum neurons were initially voltage clamped in whole-cell configuration using borosilicate glass pipettes (2-4MΩ). filled with internal solution containing (in mM): 125 K^+^-gluconate, 4 NaCl, 10 HEPES, 4 MgATP, 0.3 Na-GTP, and 10 Na-phosphocreatine (pH 7.30-7.35). Interneurons and pyramidal neurons were identified using previously established electrophysiological features (Graf et al. eNeuro 2020). Interneurons were classified by having a capacitance of less than 90 pF and an action potential half width of less than 1.5 ms. Pyramidal neurons were classified by having a capacitance of greater than 100 pF and an action potential half width of greater than 1.6 ms. Any neurons with values only meeting one of the criteria were excluded from the dataset. Following break-in to the cell, we waited ≥ 3 minutes to allow for exchange of internal solution and stabilization of membrane properties. Neurons with an access resistance of > 30MΩ or that exhibited greater than a 20% change in access resistance during the recording were not included in our datasets. After allowing the neurons to stabilize following break in, they were switched into current-clamp mode for the collection of the data presented in Figure 2. Cells that fired prior to depolarization steps were excluded from the dataset.

### Fiber photometry

Fiber photometry recordings were performed as previously described82. Briefly, an optic fiber was attached to the implanted fiber by a ferrule sleeve, then GCaMP6f was stimulated by two LEDs, a 531-Hz sinusoidal light (Thorlabs M470F3), bandpass filtered at 470 ± 20nm, and a 211-Hz sinusoidal light (Thorlabs M405FP1), bandpass filtered at 405 ± 10nm. (Filter cube: Doric FMC4; LED driver: DC4104). The 470 nm signal evokes Ca2+-dependent emission, while the 405 nm signal evokes Ca2+-independent isosbestic control emission. Prior to recording, a 180s period of GCaMP6s excitation with both light channels was used to remove the majority of baseline drift. Laser intensity at the optic fiber tip was adjusted to 50 μW before each day of recording. GCaMP6f fluorescent signal was isolated by bandpass filtering (525 ± 25nm), transduced by a femtowatt silicon photoreceiver (Newport 2151), and recorded by a real-time processor (TDT RZ5P). The envelopes of 531 Hz and 211 Hz signals were extracted in real time by the TDT program Synapse at a sampling rate of 1017.25 Hz. The 565nm LED was used for aIC photostimulation, with light intensity adjusted to 2.5µW at the optical fiber tip, while GCaMP signals were simultaneously recorded from the same patch cable at CLA.

Fiber recordings were analyzed using custom MATLAB scripts available on Github. The isosbestic signal was subtracted from the calcium-dependent signal, then baseline drift due to slow photobleaching artifacts was corrected by fitting a double exponential curve to the raw trace. The baseline-corrected signal was low-pass filtered using a Butterworth filter with cutoff set at 10 Hz. The filtered signal was then normalized by subtracting the median signal value and dividing by the absolute median value of the isosbestic signal. The correct and normalized photometry trace was z-scored relative to the mean and standard deviation of the signal over the entire trial. The mean z-score during comparable periods of time determined by the stimulus duration (when applicable) were compared using paired t-tests.

### One-photon endoscope imaging

For in vivo Ca2+ imaging experiments, we inject AAV5-CaMKIIα-GCAMP6f (1.31×1013 VG/ml, 1:3 dilution) to ensure optimal expression within the DG to ensure accurate sparse labeling for ideal imaging conditions. We then injected unilaterally with 400 nl of virus into the CLA area using a Hamilton syringe with blunted needle. After waiting at least three weeks to allow for viral spread, we implanted a microendo-scopic GRIN lens (1mm diameter x 6mm in length) 100 microns above the previously injected CLA target using the previously described protocol (Resendez et al., 2016). Briefly, using a trephine drill bit, a circular portion of the skull was removed from above the viral injection site and exposed tissue was washed with saline. The GRIN lens was then slowly lowered (100µm/min) into place with a stable stereotaxic holder attachment. Superglue was used to fix the lens in place and prevent movement during imaging, and a dental cement wall was built on the skull surrounding the lens with 2-3 anchor screws inserted into surrounding areas of the skull for support. The lens was then covered with silicone elastomer sealant (Kwik-Cast) for 1-2 weeks before checking for visibly fluorescent cells. Once the best focus was obtained, a baseplate (Inscopix) was mounted with superglue on top of the skullcap, surrounding the lens at the optimal focal distance for imaging.

nVista acquisition software (Inscopix) was used to acquire images of fluorescence dynamics at 20 frames per second. A webcame programmed to trigger the beginning of the imaging session with recording of behavior simultaneously using Ethovision software. Then, calcium imaging data were down-sampled temporally (2x temporal bin) and spatially (2x spatial bin), concatenated together, and rigid motion correction was applied. After preprocessing, putative single neuron activity was segmented using Constrained Non-negative Matrix Factorization for Endoscopic data (CNMFe) according to established criteria using custom MATLAB (MathWorks, Natick, MA, USA) scripts and sorted manually by a blind observer according to established criteria (Resendez et al., 2016; Zhou et al., 2018). Background components were estimated using the “ring” model, which was chosen after thorough parametrization to limit the presence of non-neuronal signals in our dataset.

Following sorting, individual putative single neurons were tracked between NS/FS or Sline/U50 conditions using CellReg (Sheintuch et al., 2017). For each mouse the spatial correlation registration threshold performed best, and optimal correlation thresholds were determined by the algorithm. Final registration utilized the probabilistic model for all mice.

In order to classify cells as selective for activated or inhibited during feeding onset under different recording conditions, we compare event traces during the −20 to 20 seconds before and after HPD feeding onset scored mannually for each day. A critical sigma >0.75 was chosen to determine whether a cell was significantly more active or inhibited on a given day.

### Statistical analyses

All summary data are expressed as mean ± SEM. Statistical significance was taken as *p < 0.05, **p < 0.01, ***p < 0.001, ****p < 0.0001, as determined by two-tailed Student’s t-test (paired and unpaired), two-way repeated measure analysis of variance (ANOVA) followed by Tukey’s multiple comparison test, two-tailed Wilcoxon matched-pairs signed rank test, and Friedman repeated measures ANOVA followed by Dunn’s multiple comparison test. Statistical analyses were performed in GraphPad Prism 9. For sequencing experiments, analyses were performed as described in the corresponding methods section.

## ACKNOWLEDGMENTS

We thank all of Bruchas lab for their helpful insights and discussions with manuscript. We thank Lin Tian (UCD) for providing klight viruses.

## DATA AVAILABILITY

The data supporting this study’s findings are available upon request from the authors.

## SUPPORTING INFORMATION

Additional supporting information may be found in the online version of the article.

## SUPPLEMENTARY FIGURES

**Supplementary Figure S1.**
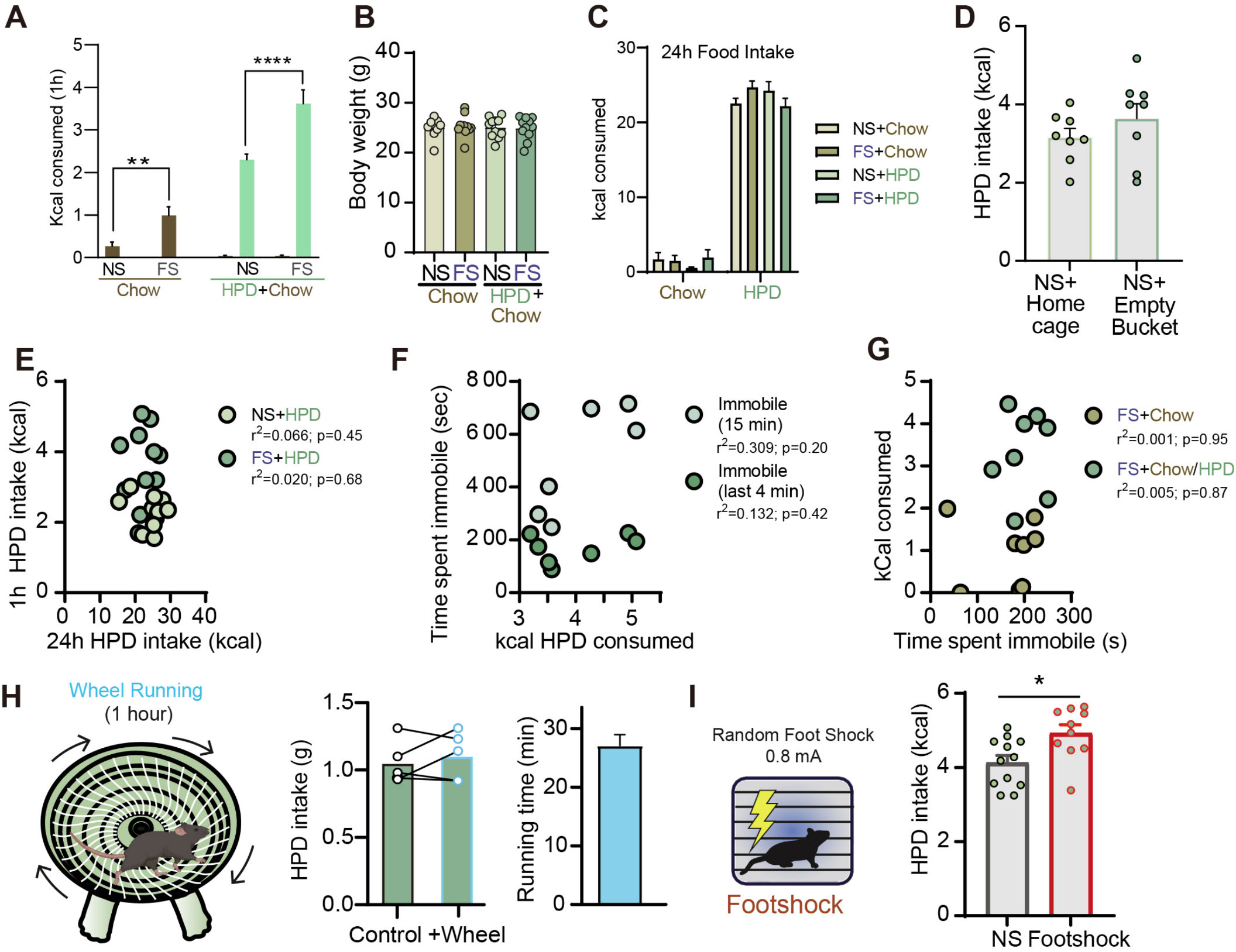
Behavioral controls for stress-associated feeding. (**A–D**) One-hour caloric intake with chow alone or HPD plus chow (**A**), body weight (**B**; n = 10 mice per group), 24-h intake (**C**), and comparison of home-cage versus empty-beaker NS procedures (**D**; n = 8). (**E–G**) Associations of intake and forced-swim immobility, with original r^2^ and P values. (**H**) Wheel-running control. (**I**) Intake after NS or random footshock (0.8 mA; n = 10). Original control data and annotations are retained; bars with errors are mean ± s.e.m. Source symbol definitions: *P < 0.05, **P < 0.01, ****P < 0.0001.

**Supplementary Figure S2.**
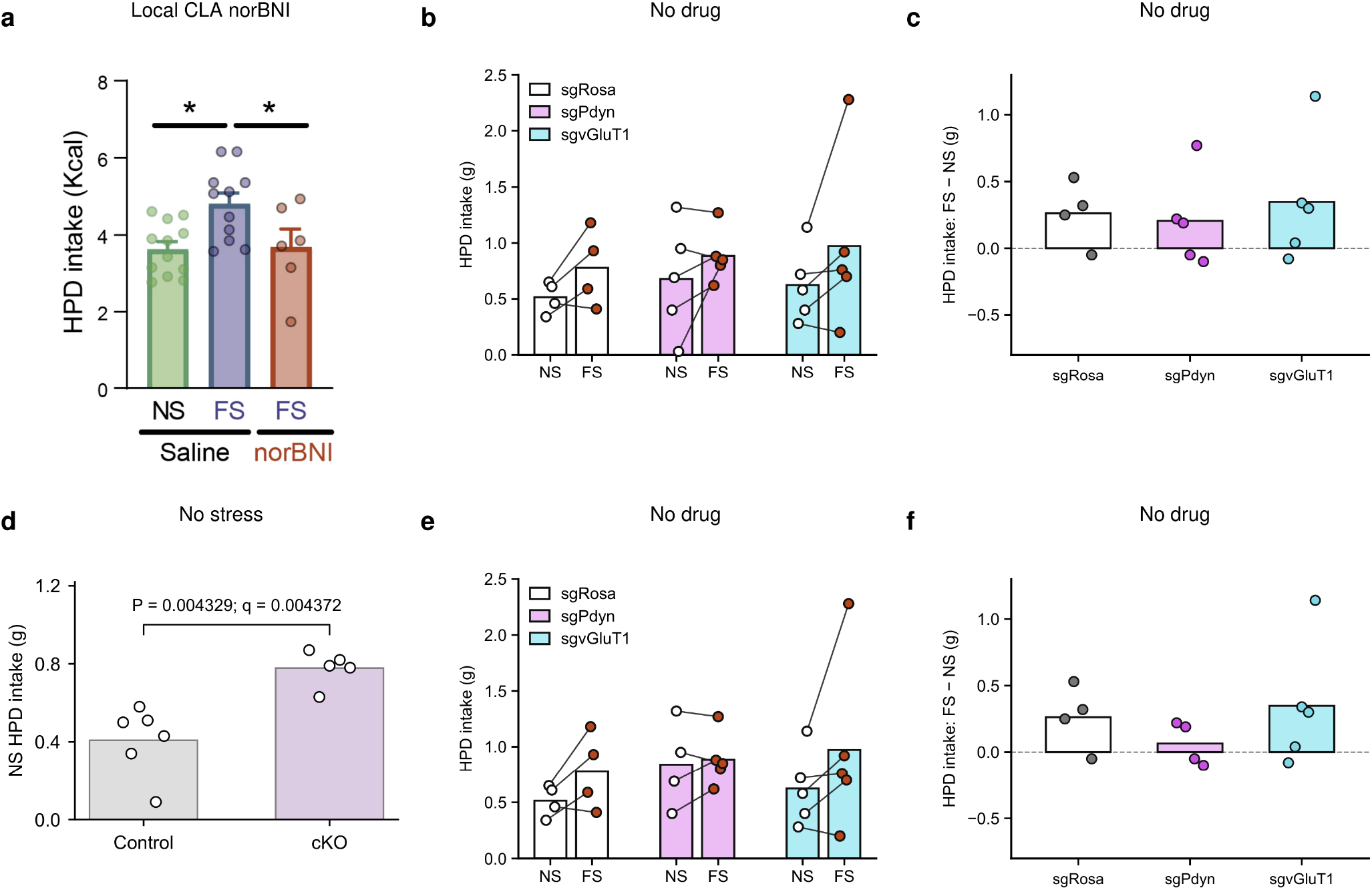
Intake after KOR blockade and Pdyn-pathway perturbations. **a**, One-hour HPD intake after intra-CLA saline under NS or FS, or norBNI under FS; source n = 6–10 mice per group, mean ± s.e.m., *P < 0.05. **b**,**c**, No-drug CRISPR intake for all recorded mice: paired NS/FS intake (**b**) and within-mouse FS−NS differences (**c**); sgRosa, sgPdyn and sgvGluT1 n = 4, 5 and 5, respectively. **d**, No-stress intake in the independent WT control (n = 6) and Pdyn cKO (n = 5) cohort. The original exact two-sided Mann– Whitney comparison gives U = 0, P = 0.004329 and q = 0.004372; q follows the original two-stage Benjamini–Krieger–Yekutieli FDR correction across the NS and FS group comparisons at Q = 1%. **e**,**f**, The same CRISPR cohort as in **b**,**c** after the author-reviewed exclusion of one sgPdyn NS session (0.03 g), selected retrospectively after inspection of intake and video. The corresponding FS value (0.80 g) remains as an unpaired observation in **e** and contributes no paired difference in **f**. NS/FS sample sizes in **e** are 4/4, 4/5 and 5/5; complete-pair sample sizes in **f** are 4, 4 and 5 (sgRosa, sgPdyn and sgvGluT1). Both inclusion views are displayed; they are not independent cohorts. In **b–f**, bars show means without error bars; points are individual mouse-session values or within-mouse differences; connecting lines identify paired observations. Intake is in grams without body-weight normalization. NS, no stress; FS, forced swim.

**Supplementary Figure S3.**
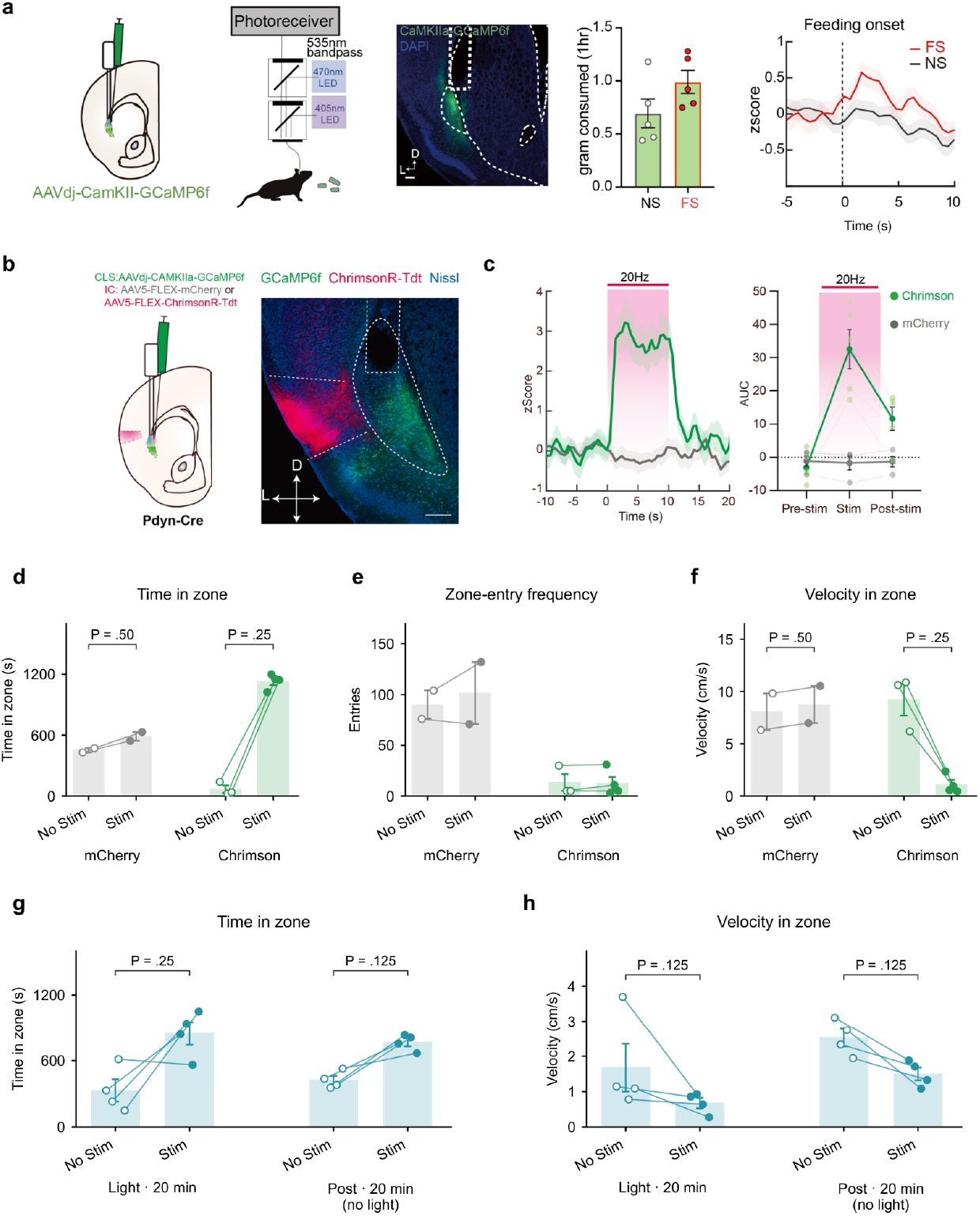
Claustral photometry and place occupancy during somatic and terminal stimulation of aIC Pdyn neurons. **a**, CaMKII-driven GCaMP6f expression in the claustrum, photometry configuration, representative expression and fiber placement, one-hour high-palatability-diet (HPD) intake, and feeding-onset-aligned fluorescence, shown from left to right (n = 5 mice). Gray/black, no stress (NS); red, forced swim (FS). Fluorescence retains its original z-score display. These are CaMKII-targeted bulk recordings, distinct from the KOR-Cre single-cell measurements in Main Figure 3. **b**, Cre-dependent ChrimsonR or mCherry in aIC Pdyn neurons, together with CaMKII-GCaMP6f in the CLA of Pdyn-Cre mice, and representative expression. **c**, Claustral fluorescence time course and pre-/during-/post-stimulation area under the curve (AUC) during 20-Hz stimulation with 10-ms pulses for 10 s. Green, ChrimsonR (n = 5 mice); gray, mCherry (n = 4). Shading marks the stimulation period. This is a representative stimulation condition; it does not isolate dynorphin from other transmitters released by aIC Pdyn axons. **d–f**, Somatic stimulation of aIC Pdyn neurons: time in zone (**d**), zone entries (**e**) and velocity (**f**). Gray, mCherry (n = 2 mice); green, Chrimson (n = 4 mice). Bars show mean ± s.e.m.; points represent individual mice and lines connect complete pairs. One Chrimson mouse has no valid No Stim observation, giving four Stim observations, three No Stim observations and three complete pairs per measure; controls have two complete pairs. **g**,**h**, Time in zone (**g**) and velocity (**h**) during terminal stimulation of aIC Pdyn neurons in the CLA, followed by a separate session without stimulation (n = 4 mice in both sessions). Each session lasted 20 min. During Post, Stim denotes the previously stimulation-paired zone. Bars show mean ± s.e.m.; points represent mice and lines pair zones within each session. Bracket P values in **d**,**f–h** are unadjusted two-sided exact within-mouse sign-flip tests. Holm-adjusted P: **d**, 0.50 for both groups; **f**, 0.75 for both groups; **g**,**h**, 0.50 in each session. Corrections retain the original four-test velocity and 2025 phase families; the two new PG duration tests form a separate family. Sensitivity t-tests and model-based 95% confidence intervals are tabulated separately. Occupancy differences accompany slowing and do not establish reward-specific effects.

**Supplementary Figure S4.**
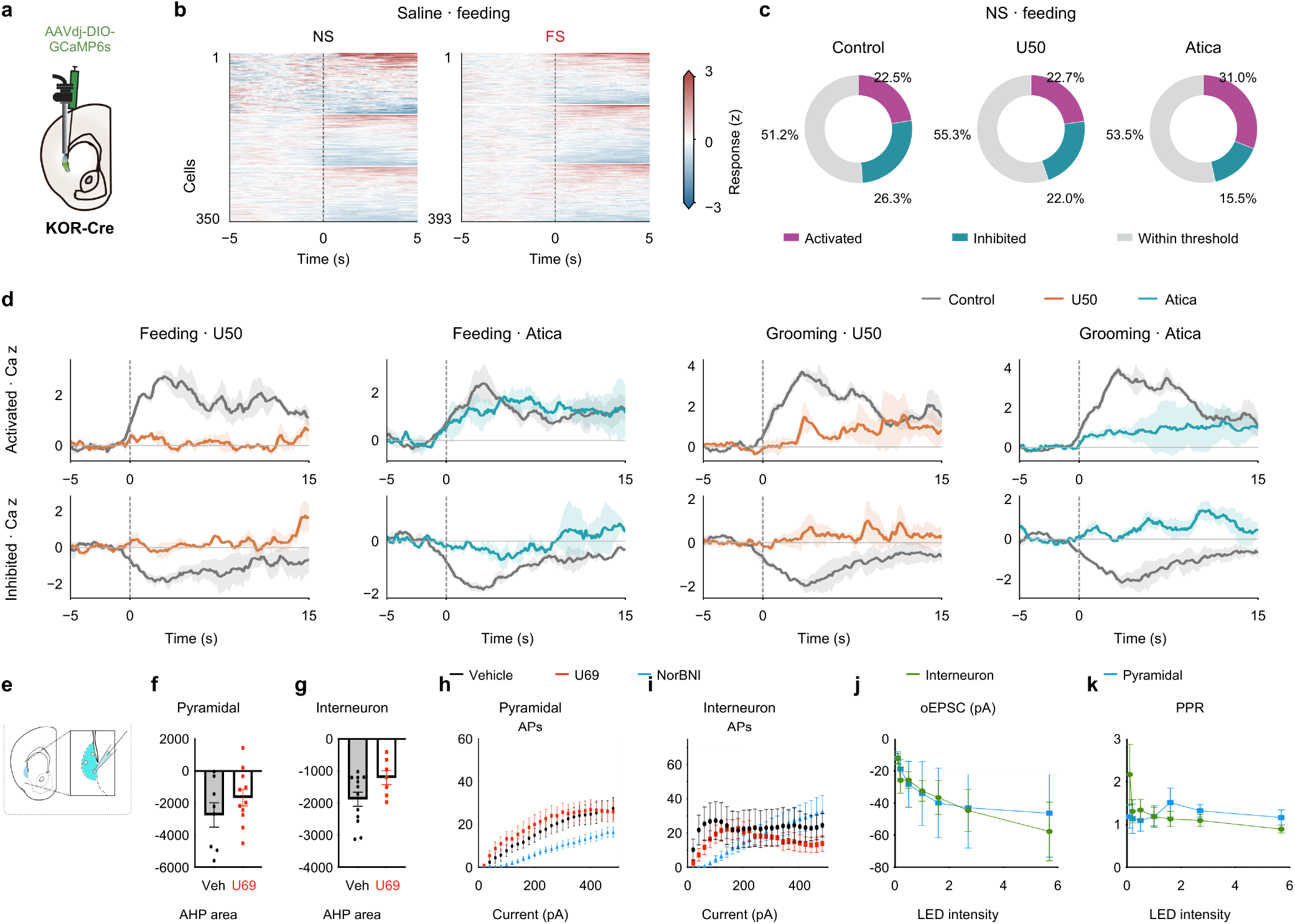
Claustral calcium responses and whole-cell electrophysiology. **a**, Author-provided KOR-Cre one-photon imaging schematic (AAVdj-DIO-GCaMP6s). **b**, Feeding-onset responses under saline NS and FS from three mice, using the common processing in Fig. 3**l–o**. Heatmaps show 350 NS and 393 FS cell-session rows, grouped by mouse and sorted within recording; cross-day cell identity is not assumed. Responses use the pooled within-event baseline SD. Colors share limits of −3 to +3 z and saturate beyond them. **c**, Original NS feeding-response classes under pooled No Drug/Saline Control, U50 or aticaprant (Atica). Fractions use 654, 436 and 155 cell-session rows from 3, 3 and 1 mouse, respectively. Scores above +1 or below −1 define descriptive activated/inhibited classes; other cells are within threshold. This score uses the baseline SD of the trial-averaged trace and differs from the common-processing sign metric in Fig. 3. **d**, Matched candidate-cell drug responses with classes fixed from Control. Feeding comparisons include 70 activated/67 inhibited cells under U50 (3 mice) and 49/63 under Atica (2 mice); grooming comparisons include 68 activated/76 inhibited cells under U50 and 75/67 under Atica (3 mice each). Curves show equal-mouse means ± between-mouse SEM after within-mouse cell averaging. Atica feeding includes KG2’s single annotated bout and KG3’s two bouts; KG2 does not enter the repeated-bout classification in **c**. The original drug plots are retained. Geometric matching and Control-based selection do not establish ensemble switching or cellwise significance. **e**, Whole-cell recording schematic. **f**,**g**, Afterhyperpolarization (AHP) area in source-defined pyramidal neurons (**f**; Vehicle/U69, 8/11 recorded-cell observations) and interneurons (**g**; 12/7). Bars and error bars retain the author figure; individual observations are shown. The saved two-sided unpaired t test for **g** gives t(17) = 2.013, P = 0.0602. No corresponding saved test was found for **f. h**,**i**, Action potentials evoked by 20–480-pA current steps. **h** was reconstructed from the supplied Prism table as means ± SEM to include the U69 series. Pyramidal-cell n at 20–80 pA is 14/13/8 and at 100–480 pA is 15/13/8 for Vehicle/U69/NorBNI, respectively. **i** retains the author plot (12/10/11 included interneuron columns). Source exclusion flags were retained; no new exclusions were made. **j**,**k**, Optically evoked EPSC amplitude (**j**) and paired-pulse ratio (**k**) versus the source LED-intensity settings; interneurons, n = 8; pyramidal neurons, n = 4. The saved two-sided unpaired t test at the highest PPR setting (5.7) gives t(10) = 1.508, P = 0.163. These are cell-level records. The supplied Prism file does not identify individual mice or map recorded cells to mice. Pyramidal/interneuron designations are retained and do not establish KOR-Cre/vGAT identities. AHP units, LED calibration and drug concentrations require confirmation.

**Supplementary Figure S5.**
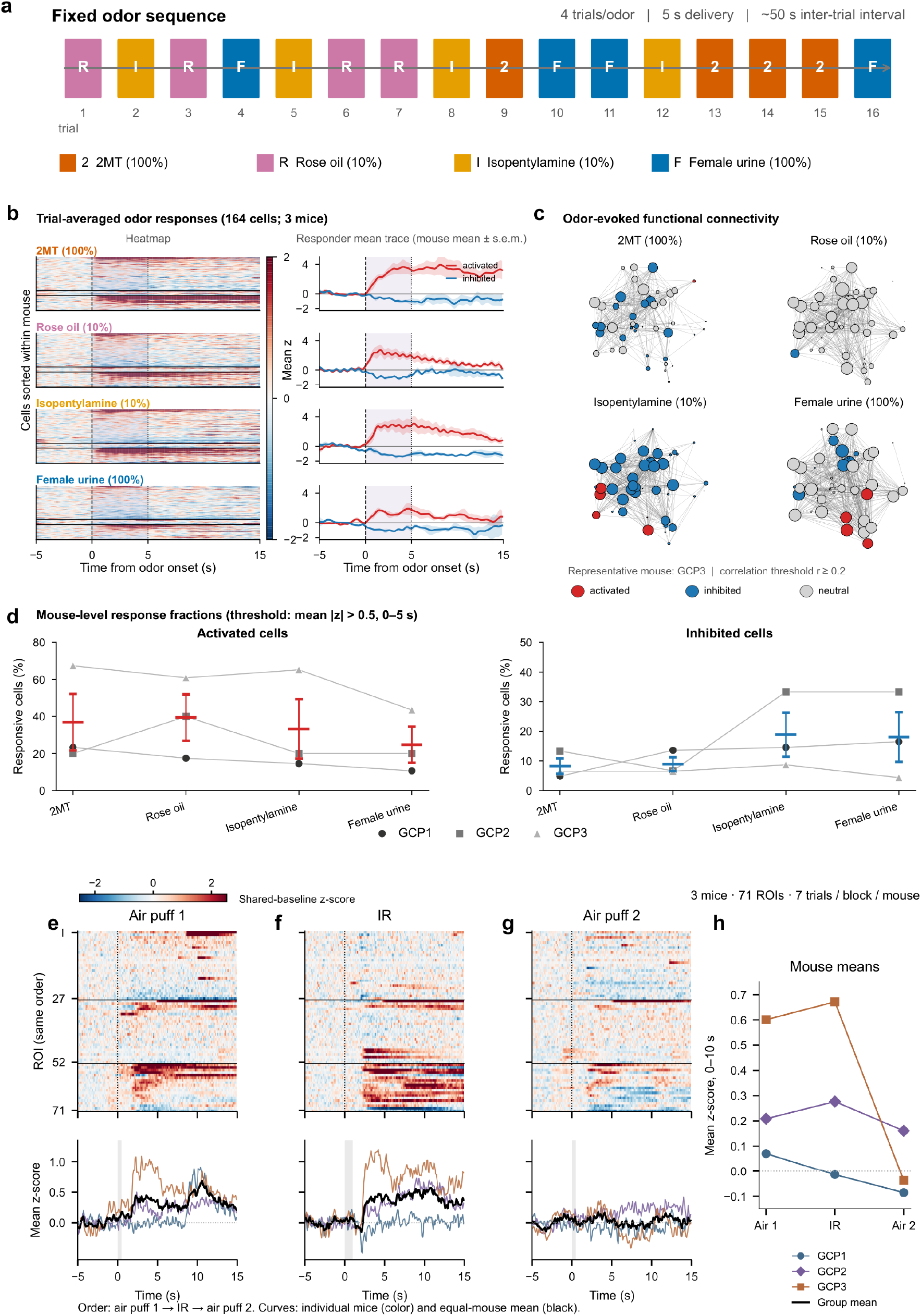
Claustral vGAT calcium responses to olfactory, air-puff and infrared stimuli. **a**, Fixed sequence of four olfactory stimuli (2MT, rose oil, isopentylamine and female urine), with four 5-s deliveries per odor and approximately 50 s between deliveries. **b**, Trial-averaged olfactory responses from 164 Suite2p-classified cells in three mice (GCP1, 103 cells; GCP2, 15; GCP3, 46). Heatmaps are sorted by the response within each mouse separately for each odor. Red and blue traces show the equal-mouse mean ± s.e.m. for cells with a mean 0–5-s z-score >0.5 or <−0.5, respectively. These are descriptive response categories, not cellwise significance tests. Fluorescence was corrected as F − 0.7Fneu; trials were averaged before normalization to the −5 to 0-s baseline and Gaussian smoothing (sigma, 1.5 samples). The supplied GCP1 alignment retains its 1,020-s imaging-duration assumption; the corresponding voltage record ends at 900 s, and acquisition-level confirmation remains pending. **c**, Representative odor-associated coactivity networks from GCP3, retained from the supplied analysis (source-labelled threshold, 0.2). Red, blue and gray retain the source positive, negative and neutral response labels. The four exports contain 46, 46, 45 and 46 nodes for 2MT, rose oil, isopentylamine and female urine, respectively. These inherited maps are descriptive illustrations, not a matched-ROI or across-mouse connectivity test. **d**, Positive- and negative-response fractions for each mouse across odors, using the thresholds in **b**; colored summary marks indicate mean ± s.e.m. (n = 3 mice). **e–g**, Responses to the first air-puff block (**e**), infrared (IR) stimulation (**f**) and the second air-puff block (**g**). Upper panels show the same 71 approved ROIs (27, 25 and 19 from GCP1, GCP2 and GCP3), sorted within mouse by the first air-puff response; horizontal lines separate mice. Lower panels show each mouse mean (colored) and the equal-mouse mean (black). Gray shading indicates approximate stimulation duration (air puff, 0.5 s; IR, 1 s). Seven valid trials were balanced chronologically per mouse per block. After F − 0.7Fneu correction, each trial was centered to its −5 to 0-s mean and divided by the same ROI’s pooled pre-event residual standard deviation across all three blocks. **h**, Mean 0–10-s response per mouse across the fixed air puff → IR → air puff sequence (n = 3 mice); connected points identify the same mouse. Responses are descriptive; no new hypothesis test was added. The fixed sequence does not distinguish prior IR exposure from elapsed time or habituation. Odor and air-puff/IR panels use different normalization procedures and ROI sets, so their z-score magnitudes are not directly comparable.

## REFERENCES

Abraham, A. D. et al. Estrogen regulation of GRK2 inactivates kappa opioid receptor signaling mediating analgesia, but not aversion. Journal of Neuroscience 38, 8031–8043 (2018). 10.1523/JNEUROSCI.0653-18.2018

Adam, T. C. & Epel, E. S. Stress, eating and the reward system. Physiology & Behavior 91, 449–458 (2007).

Anversa, R. G. et al. A model of emotional stress-induced binge eating in female mice with no history of food restriction. Genes, Brain and Behavior 19, e12613 (2020).

Bolger, M., Jackson, J. & Taylor, A. M. W. Opioid receptor distribution in the claustrum–dorsal endopiriform complex. iScience 29, 116213 (2026). 10.1016/j.isci.2026.116213

Bruchas, M. R., Land, B. B. & Chavkin, C. The dynorphin/kappa opioid system as a modulator of stress-induced and pro-addictive behaviors. Brain Research 1314, 44–55 (2010).

Cambridge, V. C. et al. Neural and behavioral effects of a novel mu opioid receptor antagonist in binge-eating obese people. Biological Psychiatry 73, 887–894 (2013).

de Almeida, C. et al. The claustrum encodes and scales appetitive feeding. Neuropsychopharmacology (2026). 10.1038/s41386-026-02573-1

Dong, C., Gowrishankar, R., Jin, Y. et al. Unlocking opioid neuropeptide dynamics with genetically encoded biosensors. Nature Neuroscience 27, 1844–1857 (2024). 10.1038/s41593-024-01697-1

Gogolla, N. The insular cortex. Current Biology 27, R580–R586 (2017).

Graf, M., Nair, A., Wong, K. L. L., Tang, Y. & Augustine, G. J. (2020). Identification of Mouse Claustral Neuron Types Based on Their Intrinsic Electrical Properties. eNeuro 7(4), ENEURO.0216-20.2020. 10.1523/ENEURO.0216-20.2020

Jackson, J., Smith, J. B. & Lee, A. K. The anatomy and physiology of claustrum–cortex interactions.Annual Review of Neuroscience 43, 231–247 (2020).

Kalueff, A. V. et al. Neurobiology of rodent self-grooming and its value for translational neuroscience. Nature Reviews Neuroscience 17, 45–59 (2016). 10.1038/nrn.2015.8

Marvin, J. S. et al. An optimized fluorescent probe for visualizing glutamate neurotransmission. Nature Methods 10, 162–170 (2013).

Niu, M. et al. Claustrum mediates bidirectional and reversible control of stress-induced anxiety responses. Science Advances 8, eabi6375 (2022).

Pereira, T. D. et al. SLEAP: a deep learning system for multi-animal pose tracking. Nature Methods 19, 486–495 (2022).

Redila, V. A. & Chavkin, C. (2008). Stress-induced reinstatement of cocaine seeking is mediated by the kappa opioid system. Psychopharmacology 200(1), 59–70. 10.1007/s00213-008-1122-y

Resendez, S. L., Jennings, J. H., Ung, R. L., Namboodiri, V. M. K., Zhou, Z. C., Otis, J. M. et al. (2016). Visualization of cortical, subcortical and deep brain neural circuit dynamics during naturalistic mammalian behavior with head-mounted microscopes and chronically implanted lenses. Nature Protocols 11, 566–597. 10.1038/nprot.2016.021

Schneider, S., Lee, J. H. & Mathis, M. W. Learnable latent embeddings for joint behavioural and neural analysis. Nature 617, 360–368 (2023).

Sheintuch, L., Rubin, A., Brande-Eilat, N., Geva, N., Sadeh, N., Pinchasof, O. & Ziv, Y. (2017). Tracking the Same Neurons across Multiple Days in Ca2+ Imaging Data. Cell Reports 21(4), 1102–1115. 10.1016/j.celrep.2017.10.013

Tian, L., Dong, C., Gowrishankar, R., Jin, Y., He, X., Gupta, A. et al. (2023). Unlocking opioid neuropeptide dynamics with genetically-encoded biosensors. Research Square [Preprint]. 10.21203/rs.3.rs-2871083/v1

Wang, Y.-J. et al. The claustrum-prelimbic cortex circuit through dynorphin/κ-opioid receptor signaling underlies depression-like behaviors associated with social stress etiology. Nature Communications (2023). 10.1038/s41467-023-43636-x

Zhou, P., Resendez, S. L., Rodriguez-Romaguera, J., Jimenez, J. C., Neufeld, S. Q., Giovannucci, A. et al. (2018). Efficient and accurate extraction of in vivo calcium signals from microendoscopic video data. eLife 7, e28728. 10.7554/eLife.28728

