## Supplemental Data 1 for "Insular opioid signaling and distinct claustral ensembles in hedonic feeding"

### SUPPLEMENTARY FIGURES

#### Supplementary Figure S1. Behavioral controls for stress-associated feeding.

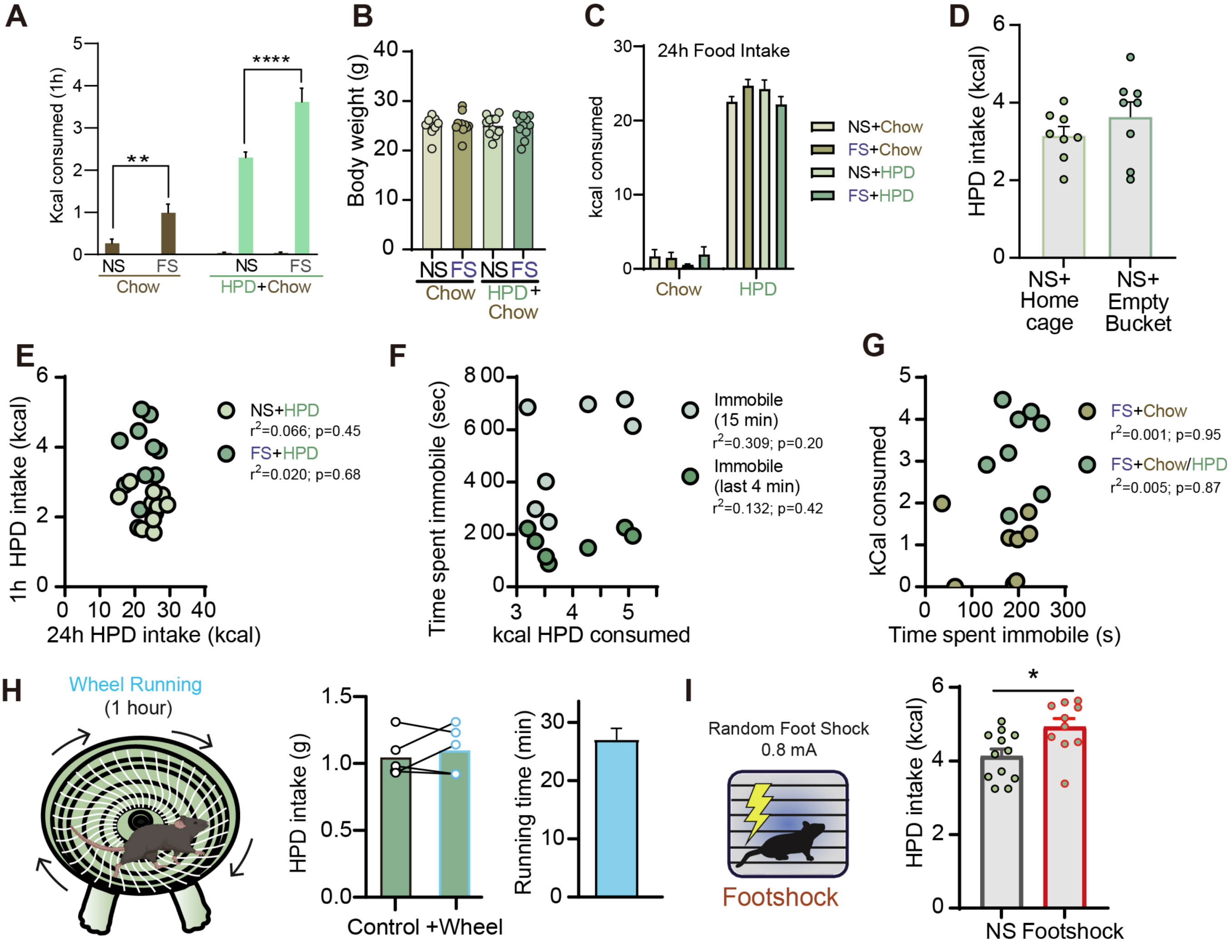


(**A–D**) One-hour caloric intake with chow alone or HPD plus chow (**A**), body weight (**B**; n = 10 mice per group), 24-h intake (**C**), and comparison of home-cage versus empty-beaker NS procedures (**D**; n = 8). (**E–G**) Associations of intake and forced-swim immobility, with original $r^2$ and P values. (**H**) Wheel-running control. (**I**) Intake after NS or random footshock (0.8 mA; n = 10). Original control data and annotations are retained; bars with errors are mean ± s.e.m. Source symbol definitions: *P < 0.05, **P < 0.01, ****P < 0.0001.

**Supplementary Figure S2. Intake after KOR blockade and Pdyn-pathway perturbations.**

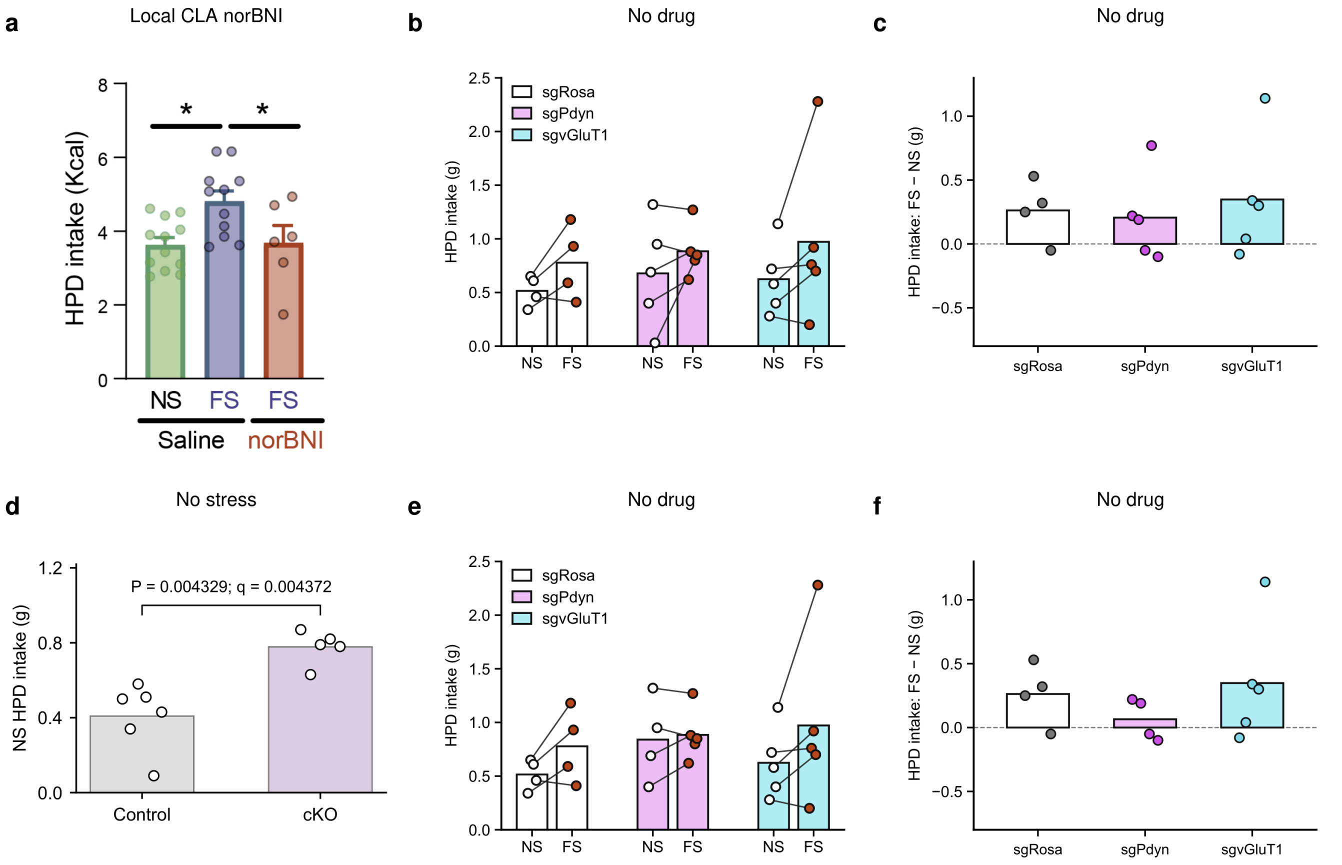


**a**, One-hour HPD intake after intra-CLA saline under NS or FS, or norBNI under FS; source n = 6–10 mice per group, mean ± s.e.m., *P < 0.05. **b,c**, No-drug CRISPR intake for all recorded mice: paired NS/FS intake (**b**) and within-mouse FS–NS differences (**c**); sgRosa, sgPdyn and sgvGluT1 n = 4, 5 and 5, respectively. **d**, No-stress intake in the independent WT control (n = 6) and Pdyn cKO (n = 5) cohort. The original exact two-sided Mann–Whitney comparison gives U = 0, P = 0.004329 and q = 0.004372; q follows the original two-stage Benjamini–Krieger–Yekutieli FDR correction across the NS and FS group comparisons at Q = 1%. **e,f**, The same CRISPR cohort as in **b,c** after the author-reviewed exclusion of one sgPdyn NS session (0.03 g), selected retrospectively after inspection of intake and video. The corresponding FS value (0.80 g) remains as an unpaired observation in **e** and contributes no paired difference in **f**. NS/FS sample sizes in **e** are 4/4, 4/5 and 5/5; complete-pair sample sizes in **f** are 4, 4 and 5 (sgRosa, sgPdyn and sgvGluT1). Both inclusion views are displayed; they are not independent cohorts. In **b–f**, bars show means without error bars; points are individual mouse-session values or within-mouse differences; connecting lines identify paired observations. Intake is in grams without body-weight normalization. NS, no stress; FS, forced swim.

**Supplementary Figure S3. Claustral photometry and place occupancy during somatic and terminal stimulation of aIC Pdyn neurons.**

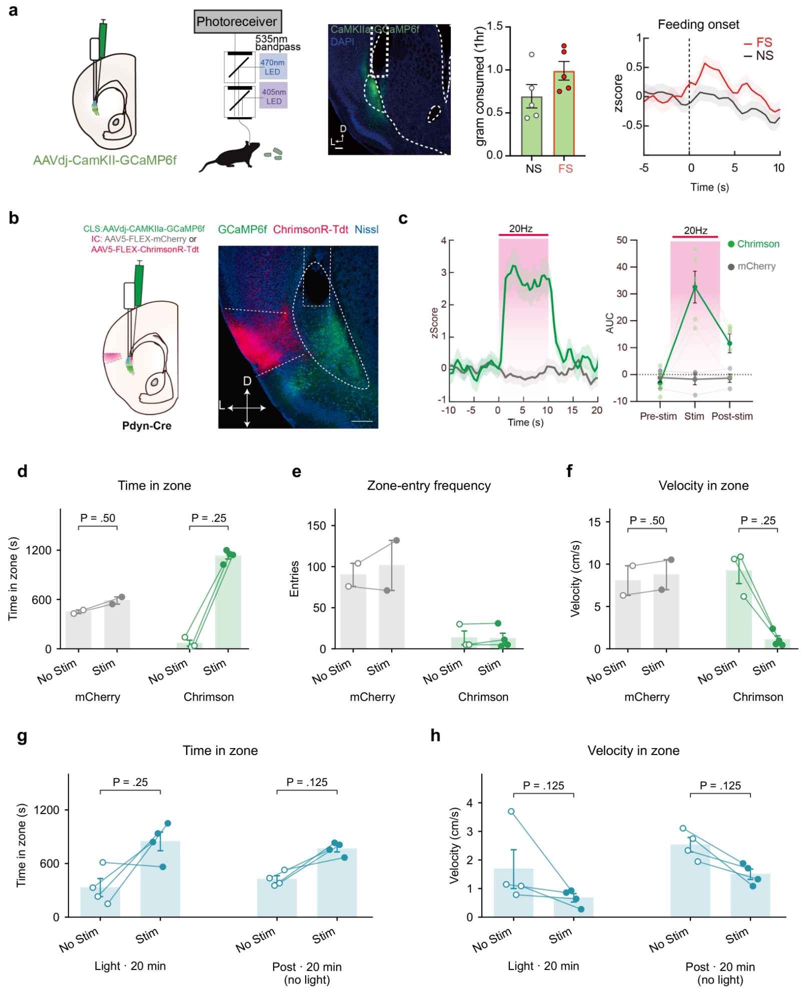


**a**, CaMKII-driven GCaMP6f expression in the claustrum, photometry configuration, representative expression and fiber placement, one-hour high-palatability-diet (HPD) intake, and feeding-onset-aligned fluorescence, shown from left to right (n = 5 mice). Gray/black, no stress (NS); red, forced swim (FS). Fluorescence retains its original z-score display. These are CaMKII-targeted bulk recordings, distinct from the KOR-Cre single-cell measurements in Main Figure 3. **b**, Cre-dependent ChrimsonR or mCherry in aIC Pdyn neurons, together with CaMKII-GCaMP6f in the CLA of Pdyn-Cre mice, and representative expression. **c**, Claustral fluorescence time course and pre-/during-/post-stimulation area under the curve (AUC) during 20-Hz stimulation with 10-ms pulses for 10 s. Green, ChrimsonR (n = 5 mice); gray, mCherry (n = 4). Shading marks the stimulation period. This is a representative stimulation condition; it does not isolate dynorphin from other transmitters released by aIC Pdyn axons. **d–f**, Somatic stimulation of aIC Pdyn neurons: time in zone (**d**), zone entries (**e**) and velocity (**f**). Gray, mCherry (n = 2 mice); green, Chrimson (n = 4 mice). Bars show mean ± s.e.m.; points represent individual mice and lines connect complete pairs. One Chrimson mouse has no valid No Stim observation, giving four Stim observations, three No Stim observations and three complete pairs per measure; controls have two complete pairs. **g,h**, Time in zone (**g**) and velocity (**h**) during terminal stimulation of aIC Pdyn neurons in the CLA, followed by a separate session without stimulation (n = 4 mice in both sessions). Each session lasted 20 min. During Post, Stim denotes the previously stimulation-paired zone. Bars show mean ± s.e.m.; points represent mice and lines pair zones within each session. Bracket P values in **d,f–h** are unadjusted two-sided exact within-mouse sign-flip tests. Holm-adjusted P: **d**, 0.50 for both groups; **f**, 0.75 for both groups; **g,h**, 0.50 in each session. Corrections retain the original four-test velocity and 2025 phase families; the two new PG duration tests form a separate family. Sensitivity t-tests and model-based 95% confidence intervals are tabulated separately. Occupancy differences accompany slowing and do not establish reward-specific effects.

#### Supplementary Figure S4. Claustral calcium responses and whole-cell electrophysiology.

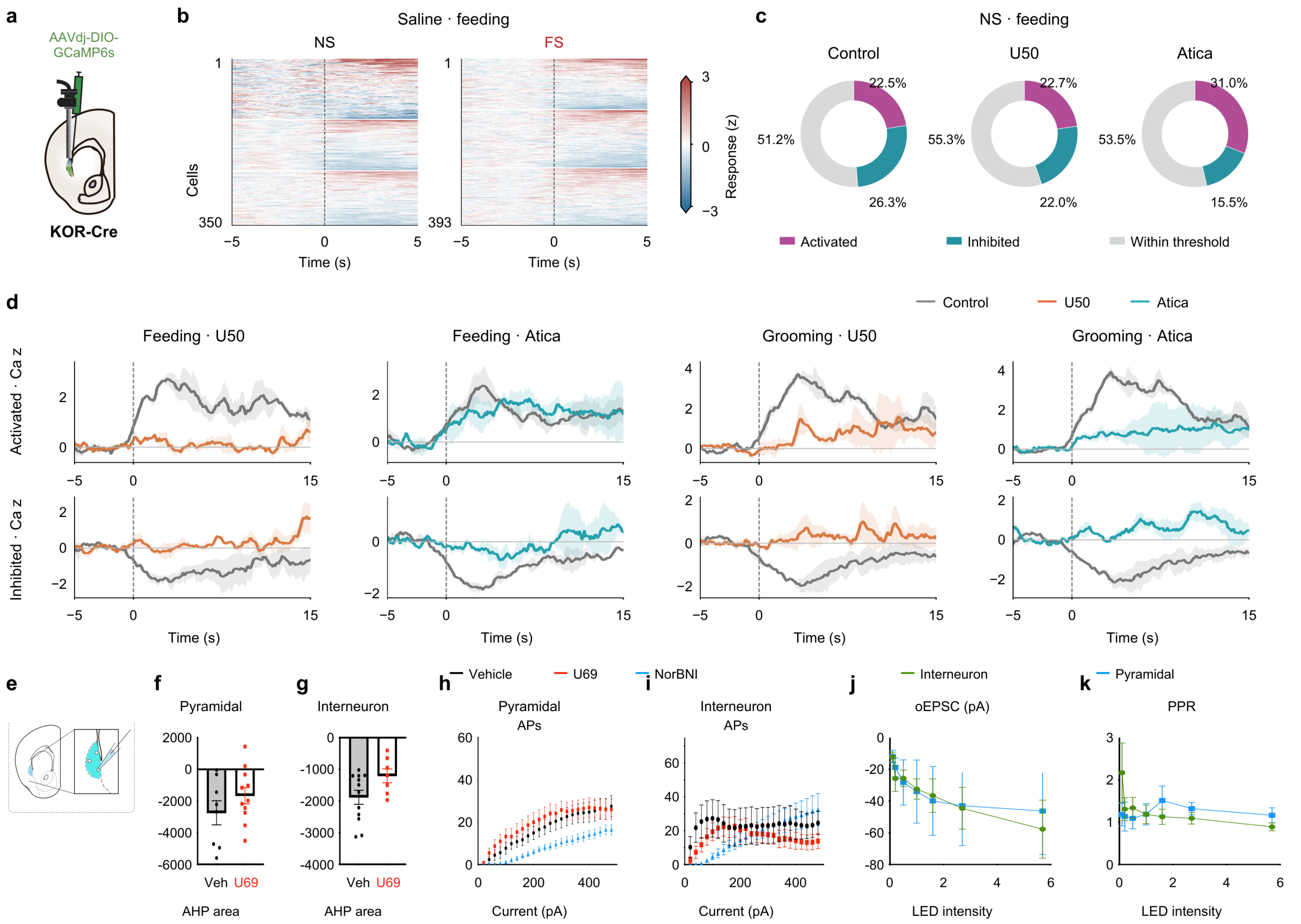


**a**, Author-provided KOR-Cre one-photon imaging schematic (AAVdj-DIO-GCaMP6s). **b**, Feeding-onset responses under saline NS and FS from three mice, using the common processing in Fig. 3**l–o**. Heatmaps show 350 NS and 393 FS cell-session rows, grouped by mouse and sorted within recording; cross-day cell identity is not assumed. Responses use the pooled within-event baseline SD. Colors share limits of −3 to +3 z and saturate beyond them. **c**, Original NS feeding-response classes under pooled No Drug/Saline Control, U50 or aticaprant (Atica). Fractions use 654, 436 and 155 cell-session rows from 3, 3 and 1 mouse, respectively. Scores above +1 or below −1 define descriptive activated/inhibited classes; other cells are within threshold. This score uses the baseline SD of the trial-averaged trace and differs from the common-processing sign metric in Fig. 3. **d**, Matched candidate-cell drug responses with classes fixed from Control. Feeding comparisons include 70 activated/67 inhibited cells under U50 (3 mice) and 49/63 under Atica (2 mice); grooming comparisons include 68 activated/76 inhibited cells under U50 and 75/67 under Atica (3 mice each). Curves show equal-mouse means ± between-mouse SEM after within-mouse cell averaging. Atica feeding includes KG2's single annotated bout and KG3's two bouts; KG2 does not enter the repeated-bout classification in **c**. The original drug plots are retained. Geometric matching and Control-based selection do not establish ensemble switching or cellwise significance. **e**, Whole-cell recording schematic. **f,g**, Afterhyperpolarization (AHP) area in source-defined pyramidal neurons (**f**; Vehicle/U69, 8/11 recorded-cell observations) and interneurons (**g**; 12/7). Bars and error bars retain the author figure; individual observations are shown. The saved two-sided unpaired t test for **g** gives t(17) = 2.013, P = 0.0602. No corresponding saved test was found for **f**. **h,i**, Action potentials evoked by 20–480-pA current steps. **h** was reconstructed from the supplied Prism table as means ± SEM to include the U69 series. Pyramidal-cell n at 20–80 pA is 14/13/8 and at 100–480 pA is 15/13/8 for Vehicle/U69/NorBNI, respectively. **i** retains the author plot (12/10/11 included interneuron columns). Source exclusion flags were retained; no new exclusions were made. **j,k**, Optically evoked EPSC amplitude (**j**) and paired-pulse ratio (**k**) versus the source LED-intensity settings; interneurons, n = 8; pyramidal neurons, n = 4. The saved two-sided unpaired t test at the highest PPR setting (5.7) gives t(10) = 1.508, P = 0.163. These are cell-level records. The supplied Prism file does not identify individual mice or map recorded cells to mice. Pyramidal/interneuron designations are retained and do not establish KOR-Cre/vGAT identities. AHP units, LED calibration and drug concentrations require confirmation.

**Supplementary Figure S5. Claustral vGAT calcium responses to olfactory, air-puff and infrared stimuli.**

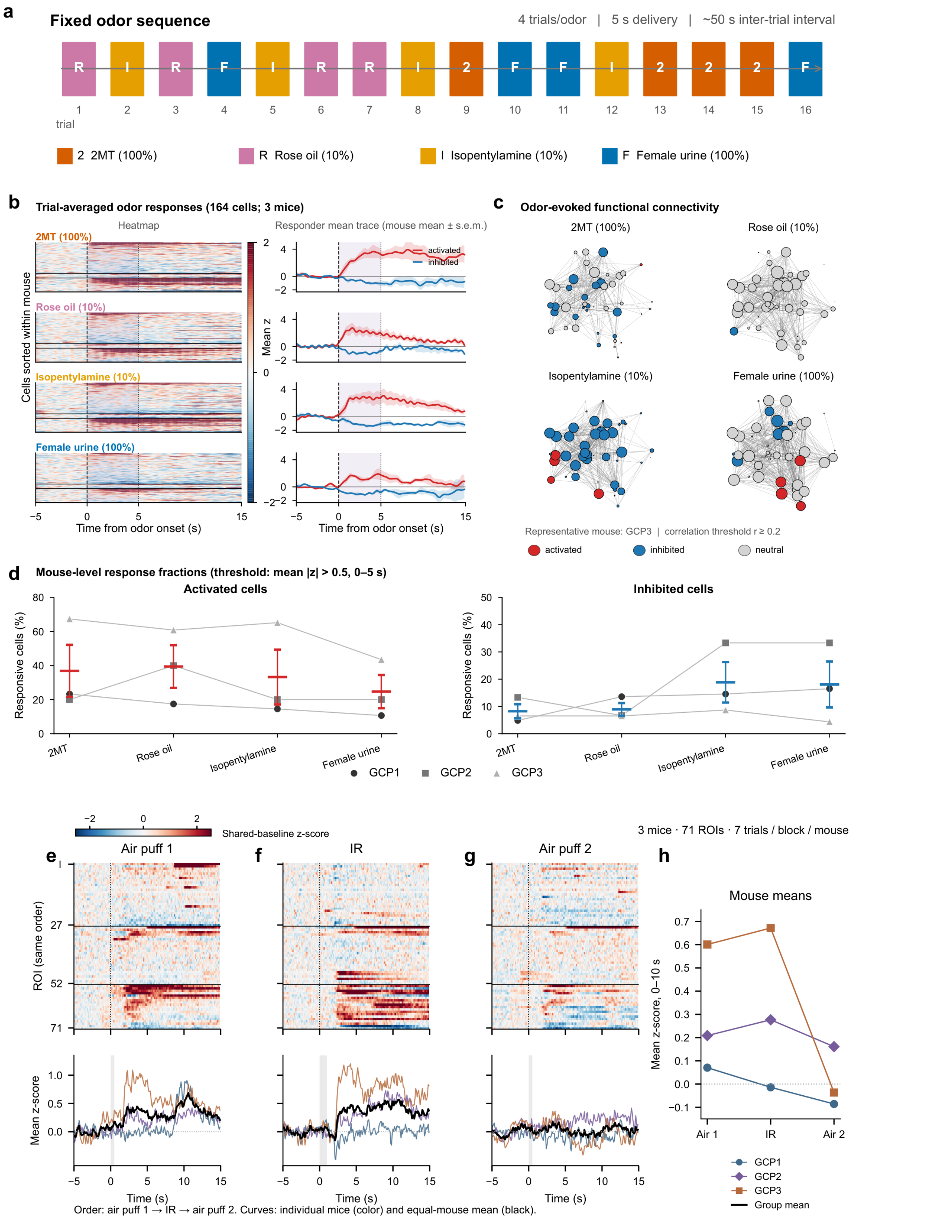


**a**, Fixed sequence of four olfactory stimuli (2MT, rose oil, isopentylamine and female urine), with four 5-s deliveries per odor and approximately 50 s between deliveries. **b**, Trial-averaged olfactory responses from 164 Suite2p-classified cells in three mice (GCP1, 103 cells; GCP2, 15; GCP3, 46).
